# Multiple routes of functional diversification of the plant BAHD acyltransferase family revealed by comparative biochemical and genomic analyses

**DOI:** 10.1101/2020.11.18.385815

**Authors:** Lars H. Kruse, Austin T. Weigle, Mohammad Irfan, Jesús Martínez-Gómez, Jason D. Chobirko, Jason E. Schaffer, Alexandra A. Bennett, Chelsea D. Specht, Joseph M. Jez, Diwakar Shukla, Gaurav D. Moghe

**Affiliations:** Plant Biology Section, School of Integrative Plant Sciences, Cornell University, Ithaca, NY, 14853, USA; L.H. Bailey Hortorium, Cornell University, Ithaca, NY, 14853, USA; Department of Chemistry, University of Illinois at Urbana-Champaign, Urbana, IL, 61801, USA; Department of Chemical and Biomolecular Engineering, University of Illinois at Urbana-Champaign, Urbana, IL, 61801, USA; Department of Molecular Biology and Genetics, Cornell University, Ithaca, NY, 14853, USA; Department of Biology, Washington University in St. Louis, St. Louis, MO, 63130, USA; Institute of Analytical Chemistry, Universität für Bodenkultur Wien, Vienna, 1190, Austria; Michael Smith Laboratories, University of British Columbia, Vancouver, British Columbia, V6T 1Z4, Canada

**Keywords:** Evolutionary biochemistry, enzyme family, comparative genomics, gene duplication, promiscuity, protein structure analysis, BAHD acyltransferase

## Abstract

Large enzyme families catalyze metabolic diversification by virtue of their ability to use diverse chemical scaffolds. How enzyme families attain such functional diversity is not clear. Here, we addressed this question using BAHD acyltransferases as a model, and identified the routes by which duplication, promiscuity and sequence changes influenced BAHD diversification. This fast-evolving family expanded drastically during land plant evolution from 1-5 copies in algae to ∼100 copies in diploid angiosperm genomes. *In vitro* characterization of fourteen BAHDs against a substrate panel and compilation of >160 published activities revealed the wide prevalence of promiscuity among BAHDs. Using phylogenetic analysis, we predicted the substrate classes that the ancestral enzymes were likely capable of using prior to land plant origins. While the anthocyanin acylation activity was fixed in BAHDs later near the origin of angiosperms, *in vitro* testing of BAHDs from non-seed plant lineages suggested that the ability to acylate anthocyanins likely existed promiscuously millions of years prior to its fixation. Motif enrichment analysis in anthocyanin-acylating BAHDs identified two motifs fixed in the largest anthocyanin acylating clade. Molecular dynamic simulations and enzyme kinetics revealed the important role of an active site tryptophan, whose bulkiness, hydrophobicity and aromaticity are critical for anthocyanin acylation. Our results thus describe the molecular processes in robust, evolvable enzymes that drive emergence of functional diversity in enzyme families.

**One sentence summary:** Using a combination of phylogenetics, biochemistry and protein structure analysis, we investigated how the BAHD acyltransferase family evolved to use a structurally diverse array of substrates.

## INTRODUCTION

Enzymes involved in plant specialized metabolism often belong to enzyme families, some of which (e.g., cytochrome P450s, lipases, acyltransferases, dioxygenases) have several hundred members in angiosperm genomes. Such enzyme families are characterized by frequent gene duplication, functional divergence, and promiscuity, all of which contribute to metabolic diversification. Due to advances in sequencing technologies, much is known today about how duplicates evolve at the genomic, epigenetic and transcriptomic levels (Ganko et al., 2007; Zou et al., 2009; Schnable et al., 2011; Moghe et al., 2014; Wang et al., 2014a); however, our understanding of how substrate preference evolves in duplicate enzymes and how promiscuity plays a role in this evolution, especially in large enzyme families, is lacking.

Promiscuity refers to the ability of an enzyme to catalyze multiple reactions, either by using different substrates (substrate promiscuity), producing multiple products from the same substrate (product promiscuity), or performing secondary reactions that cause different chemical transformations (catalytic promiscuity) (Copley, 2015). The term “enzyme promiscuity” is interpreted differently by practitioners of different fields (Copley, 2015; Khersonsky and Tawfik, 2010). In this study, substrate promiscuity is described in the context of an enzyme’s ability to use multiple substrates – a mechanistic definition popularly adopted by molecular/structural biologists (Ekroos and Sjögren, 2006; Sayou et al., 2014; Leong and Last, 2017; Kreis and Munkert, 2019; van Loo et al., 2019). Promiscuous reactions of an enzyme may get selected upon if the product directly or indirectly increases organismal fitness. Existence of such promiscuity-driven “underground metabolism” can occur via drift and contributes to the standing natural variation of metabolites (Notebaart et al., 2014).

In this study, we address the question of how gene duplication and promiscuity contribute to plant specialized metabolic diversity using the large BAHD acyltransferase family (referred to as BAHDs hereafter) as a model. Named after the four first discovered enzymes of this family – benzyl alcohol O-acetyltransferase (BEAT) (Dudareva et al., 1998), anthocyanin O-hydroxycinnamoyltransferase (AHCT) (Fujiwara et al., 1997, 1998a, 1998b), N-hydroxycinnamoyl/benzoyltransferase (HCBT) (Yang et al., 1997), and deacetylvindoline 4-O-acetyltransferase (DAT) – members of this large family catalyze the transfer of an acyl group from a coenzyme A (CoA) conjugated donor to a –OH or –NH_2_ group on an acceptor (D’Auria, 2006). The ease of heterologous protein expression in *Escherichia coli*, the intronless nature of many BAHD genes, their ability to use structurally diverse substrates, and availability of functional data from multiple species make BAHDs an attractive model system. BAHDs play important roles in the biosynthesis of several phenylpropanoids, amides, volatile esters, terpenoids, alkaloids, anthocyanins, flavonoids, and acylsugars (D’Auria, 2006; Tuominen et al., 2011). Although >160 members of this family have been experimentally characterized across the plant kingdom, transfer of known functions using sequence similarity to these characterized enzymes is difficult owing to their rapid sequence divergence, substrate promiscuity and functional divergence. For example, the 4-5 acylsugar acyltransferases involved in acylsugar biosynthesis in Solanaceae trichomes are BAHDs (Moghe et al., 2017; Leong et al., 2020) but are only 40-50% identical, and yet all of them use sucrose/acylated sucrose as substrates. In contrast, a single amino acid change is sufficient to convert a BAHD from preferentially using phenylpropanoid substrates to using amine-containing substrates (Levsh et al., 2016).

Previous studies on BAHDs have also revealed existence of substrate promiscuity (Aharoni et al., 2000; Eudes et al., 2016b; Levsh et al., 2016; Moghe et al., 2017; Chiang et al., 2018a), but compared to the known number of BAHD substrates, the extent of our knowledge about BAHD promiscuity is still limited. Furthermore, it is unclear whether BAHD family functional diversity is a result of multiple rounds of neo- functionalization – where completely new activities emerged – versus sub- functionalization of already present promiscuous side-reactions in the common ancestor of different BAHD clades.

The BAHD family is speculated to be related to the carnitine acyltransferases involved in fatty acid metabolism (St Pierre and De Luca, 2000; D’Auria, 2006), but their evolution across plants has not been studied. We were interested in characterizing evolution of the *capability*/*potential* of BAHDs to use different substrate classes rather than their *actual in vivo* substrates, since the inherent capability of an enzyme can be a starting point for selection to act and fix diversified enzyme activities in different *in vivo* contexts. Although it may be argued that BAHDs might use any substrates with hydroxyl or amine groups, previous results provide clear evidence of specialization (D’Auria, 2006), and it is unclear how these specializations emerged. We first characterized the known substrate space of extant characterized BAHDs and used it as a template in the context of BAHD phylogeny to delineate the putative ancestral substrate space. Prediction of the ancestral space further helped us identify multiple sequence and structural features important for specific functions. Overall, this study provides a template to assess functional evolution after duplication in large enzyme families, and generates resources foundational for rational prediction of BAHD function in plant genomes.

## RESULTS

### The BAHD enzyme family occupies a wide substrate space

BAHDs have been experimentally characterized across land plants (Sander and Petersen, 2011; Eudes et al., 2016b; Levsh et al., 2016; Moghe et al., 2017; Chiang et al., 2018a). To get a complete picture of the range of known BAHD substrates, we first compiled a database of 164 biochemically characterized BAHDs that used a total of 206 acceptor substrates and ∼31 acyl donors from 75 species across green plants **(Supp. Dataset 1; Supp. Dataset 2A)**. BAHDs characterized solely using other types of experimental evidence without detailing the exact substrate (i.e., gene knock-out and knock-down, gene expression analysis) were excluded from this compilation. The curated BAHD substrates were empirically classified into thirteen different types based on similarity of their chemical scaffolds and functional groups **(Supp. Table 1; Supp. Fig. 1A)** (Wang et al., 2013), and organized into a novel network representation for visualizing the already characterized BAHD substrate space **(Fig. 1A; Supp. Fig. 1B)**.

**Figure 1.**
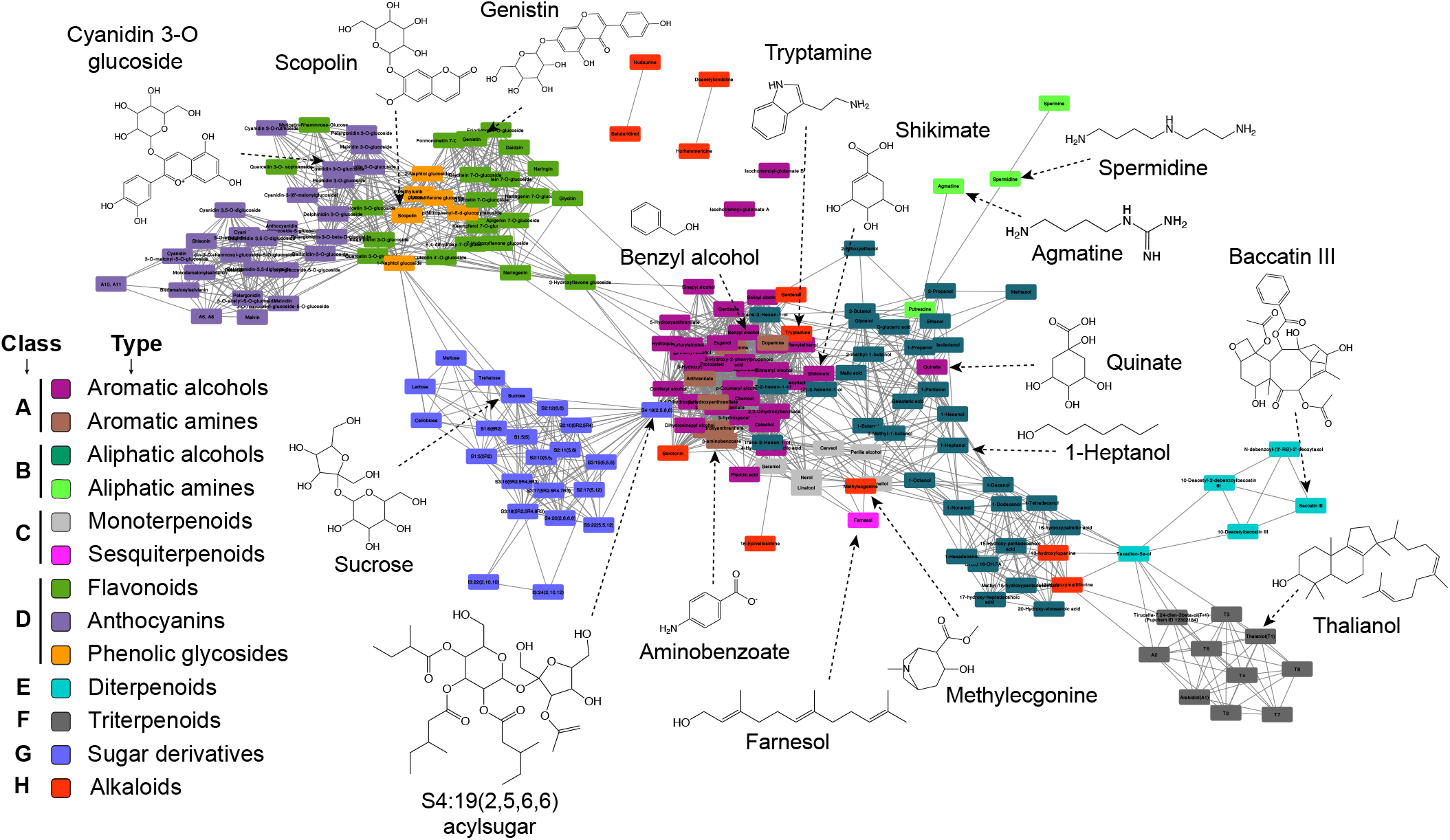
BAHDs use a structurally diverse set of substrates. Substrate similarity network of BAHD acceptor substrates (**Supp. Dataset 2B**), shown as the “Prefuse Force Directed” layout in Cytoscape. An MCS-Tanimoto similarity cutoff of 0.5 used to draw edges between two substrate nodes. Substrates are colored based on the substrate type they belong to. Substrate class H (alkaloids) is scattered accross the network and doesn’t form a distinct cluster. Exemplary structures of substrate classes are shown.

Distinct clusters obtained in this visualization revealed that BAHDs have evolved to accept at least eight different structural scaffolds **(classes A-H; Fig. 1**). A majority (109, 80%) of the characterized enzymes use substrates in four classes – class A comprising of aromatic alcohol and amines (67 enzymes, 41%), class B containing aliphatic alcohols and amines (60 enzymes, 37%), class C that includes monoterpenoids and a sesquiterpenoid (7 enzymes, 4%), and class D comprising flavonoids, anthocyanins, and phenolic glycosides (31 enzymes, 19%) **(Fig. 1; Supp. Dataset 2B)** – likely indicative of the degree of research attention on lignins, cuticular lipids, floral volatiles and pigments, respectively. BAHDs can also transform a wider array of terpenoids including mono/di/tri terpenoids in classes E and F. Some polyamines (spermine, spermidine) were more distant from aliphatic alcohols primarily due to different functional groups (-OH vs. -NH_2_), but still were classified into the same class B due to overall scaffold relatedness **(Supp. Fig. 1A; Supp. Table 1)**. Other substrates such as alkaloids (class H) and sugar derivatives (class G), represented smaller and independent classes in the network. Alkaloids themselves are a loosely defined compound type, and hence class H comprises alkaloids that could also be assigned to other classes based on scaffold similarity.

We also visualized the BAHD substrate network in the context of the larger phytochemical space represented by 7128 additional plant compounds **(Supp. Fig. 2; Supp. Dataset 3).** Since BAHDs can only use substrates containing amine or hydroxyl groups, only ∼70% of the compounds from this pool are actually available for catalysis. Nonetheless, this visualization shows that BAHDs can access a large space in the phytochemical network. Several metabolite classes (e.g., small organic acids in central metabolism, some amino acids, nitrogen bases/nucleosides, many alkaloids, peptides, polyphenols, and oligosaccharides) are structurally similar to existing BAHD substrates, thereby representing the latent catalytic potential of BAHDs that lies undetected in uncharacterized enzymes or as secondary activities of characterized enzymes.

To understand how BAHDs evolved to use so many chemical scaffolds, we focused on class A/B vs. class D substrates **(Fig. 1)**, which are structurally very distinct. Three enzymes using class A substrates (AtHCT, SmHCT, PsHCT2) were previously shown in one study to use naringenin, a flavonoid (Chiang et al., 2018a). No class D substrate using enzyme was shown to use class A/B substrates **(Supp. Dataset 2A)**. While this difference could represent true functional differentiation, it could be a result of experimenter-bias (i.e., researchers studying pigmentation may not assess lignin or volatile esters (class A/B substrates), and vice versa). Thus, BAHDs may promiscuously be able to use substrates beyond their native substrate classes – a phenomenon we refer to henceforth as “class promiscuity” to distinguish it from the ability of an enzyme to use structurally similar substrates from the same class (substrate promiscuity). Such behavior would be undetected in our collected activity database if non-native substrate classes were never tested. To determine the prevalence of class promiscuity in BAHDs, we performed extensive characterization of ten BAHDs that use class A, B (HCT/HQT- type) and class D (anthocyanin acyltransferase or AnAT-type) substrates.

### Many characterized BAHDs promiscuously use non-native substrates

We tested a total of eight previously characterized and two novel BAHD enzymes (explained in **Supp. Methods**) against an acceptor substrate array **(Supp. Dataset 4A)**, using optimal conditions described for those enzymes in previous studies. Based on the substrate networks and our activity database **(Fig. 1; Supp. Fig. 1B; Supp. Dataset 2),** we selected eleven substrates from six substrate types in classes A, B and D as acceptors for initial analysis (details on enzyme and donor substrate selection in **Supp. Methods**). Based on the results obtained using this initial substrate array, we further assayed some enzymes with additional substrates **(Supp. Dataset 4B,C)** to confirm additional hypotheses.

Of the ten enzymes, one used only a single tested substrate and six showed substrate promiscuity within the same class, which we also refer to as “specialized” below (**Supp. Dataset 4**). According to our assays and gathered literature information, 61% (105/172 enzymes) of BAHDs analyzed in this study can use more than one acceptor substrate, 21% (37/172 enzymes) use ≥5 substrates, and 7% (12/172 enzymes) can use ≥10 substrates, highlighting the considerable substrate promiscuity in the family.

Three out of ten tested BAHDs were class-promiscuous. The previously uncharacterized *Solanum lycopersicum* hydroxycinnamoyl CoA transferase (SlHCT) was highly class promiscuous **(Fig. 2; Supp. Fig. 2C; Supp. Dataset 4)**. Aromatic alcohol, amine and flavonoid use has been described for HCT/HQT-type enzymes before (Sander and Petersen, 2011; Eudes et al., 2016a; Levsh et al., 2016; Peng et al., 2016; Chiang et al., 2018a), but not anthocyanin acylation as we saw for SlHCT. No activity was found with free sucrose or glucose for any of the 10 enzymes, despite acylation on the glycoside of the anthocyanin. The liverwort enzyme MeHFT showed a similarly diverse substrate utilization pattern **(Fig. 2A)**. Note that Cucurbitacin E is not known as a substrate for any known BAHD but both SlHCT and MeHFT showed substantial activity with it. Another class-promiscuous enzyme, EcCS whose native substrate is methylecgonine (alkaloid, class H) (Schmidt et al., 2015; Eudes et al., 2016a), showed high specific activities with shikimate and quinate (class A) – two substrates that are structurally dissimilar to methylecgonine (MCS-Tanimoto = 0.47, **Supp. Dataset 2C**). These findings show that some BAHDs have specialized *in vivo* substrate profiles but have the capability to explore a large region of the phytochemical space **(Supp. Fig. 2)** through secondary activities.

**Figure 2.**
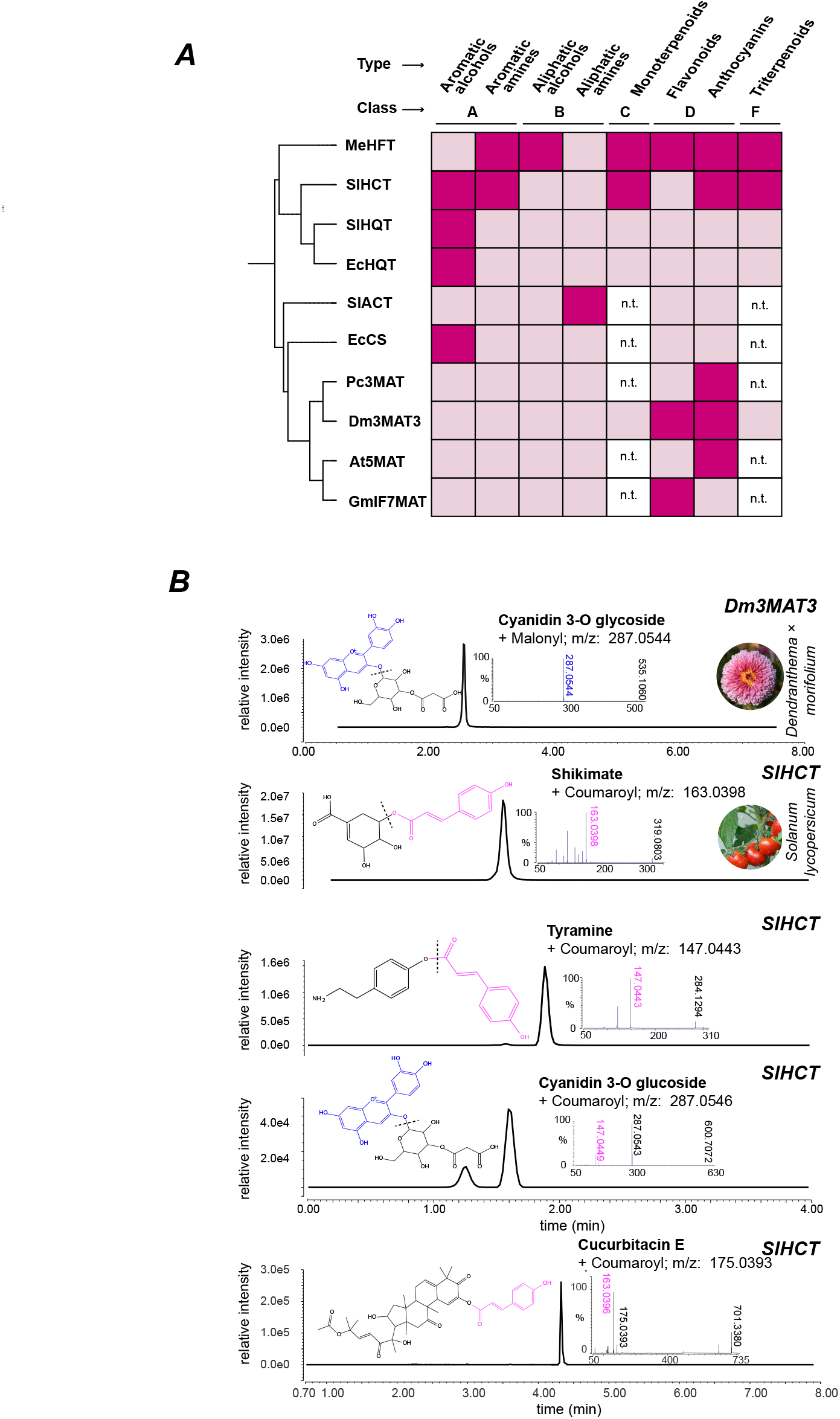
Many BAHD acyltransferases show substrate promiscuity. **(A)** Matrix of tested enzymes and substrate classes. Enzymes are ordered based on their phylogenetic relationship as determined using neighbor joining. Dark pink color indicates detected activity in the respective substrate class. See **Supp. Dataset 4** for more detailed calculations. n.t. = not tested. **(B)** Examples of extracted ion chromatograms of the quantifier ions of different enzymatic products of Dm3MAT3 and SlHCT with their preferred donor malonyl-CoA and coumaroyl-CoA, respectively. See **Supp.** Fig 3 for all other extracted ion chromatograms. Each product was measured using product-specific PRM methods (**Supp. Table 5**) and the most abundant fragment ion was used for quantification. This ion is noted in the upper right corner of each chromatogram. Structures represent the best-inference based on previously reported structures and the observed fragmentation patterns.

As opposed to the broad-specificity of enzymes using class A/B substrates, BAHDs using anthocyanins/flavonoid (class D, AnAT-type enzymes) showed substrate class usage restricted to flavonoid and anthocyanin substrates under the testing conditions **(Fig. 2; Supp. Fig. 3B; Supp. Dataset 4)**. These results suggest that enzymes that have specialized for class D substrate acylation underwent adaptations constraining them from using class A/B substrates. Since we tested AnATs using malonyl-CoA donor and class A/B-utilizing enzymes with coumaroyl- and feruloyl-CoA, we also tested whether these enzymes can acylate aromatic alcohols and anthocyanins respectively, using reciprocal donors. In both cases, no product formation was detected **(Supp. Dataset 4)**.

Through these assays, we found a previously undocumented scale of BAHD substrate and class promiscuity. Class promiscuity was restricted to enzymes using class A/B substrates (SlHCT, MeHFT, EcCS) **(Supp. Dataset 4)** and not class D substrates. We hypothesized that if class A/B represented the ancestral substrate space and if more class A/B utilizing enzymes showed class-promiscuous behavior, it would suggest that the ability to use class D substrates already existed as a promiscuous activity in ancestral enzymes. To test this hypothesis, we used phylogenetic analysis to predict the ancestral state of the BAHD family and determine patterns of diversification in the plant kingdom.

### The BAHD family expansion started in the ancestor of land plants

We first identified BAHDs from 52 sequenced plant genomes using the acyltransferase domain model PF02458. BAHDs were detected in 1-5 copies in multiple green algal genomes; however, angiosperm and gymnosperm genomes contain dozens to hundreds of copies, with non-seed plants showing intermediate BAHD counts **(Fig. 3A,C)**. Ancestral state reconstruction of BAHD counts normalized to the total gene count in each genome, using a Bounded Brownian Motion model (Boucher and Démery, 2016) **(Supp. Methods),** revealed that the relative BAHD gene content in species’ genomes began to increase prior to land plant origins **(Fig. 3A,B),** increased gradually until the emergence of seed plants, and generally stabilized in angiosperms with some lineage-specific exceptions **(Fig. 3A)**. We thus sought to determine the ancestral state of BAHD activities in the last common ancestor of land plants (where BAHDs had already started to expand) and the ancestral node of charophytes and land plants, prior to BAHD expansion.

**Figure 3.**
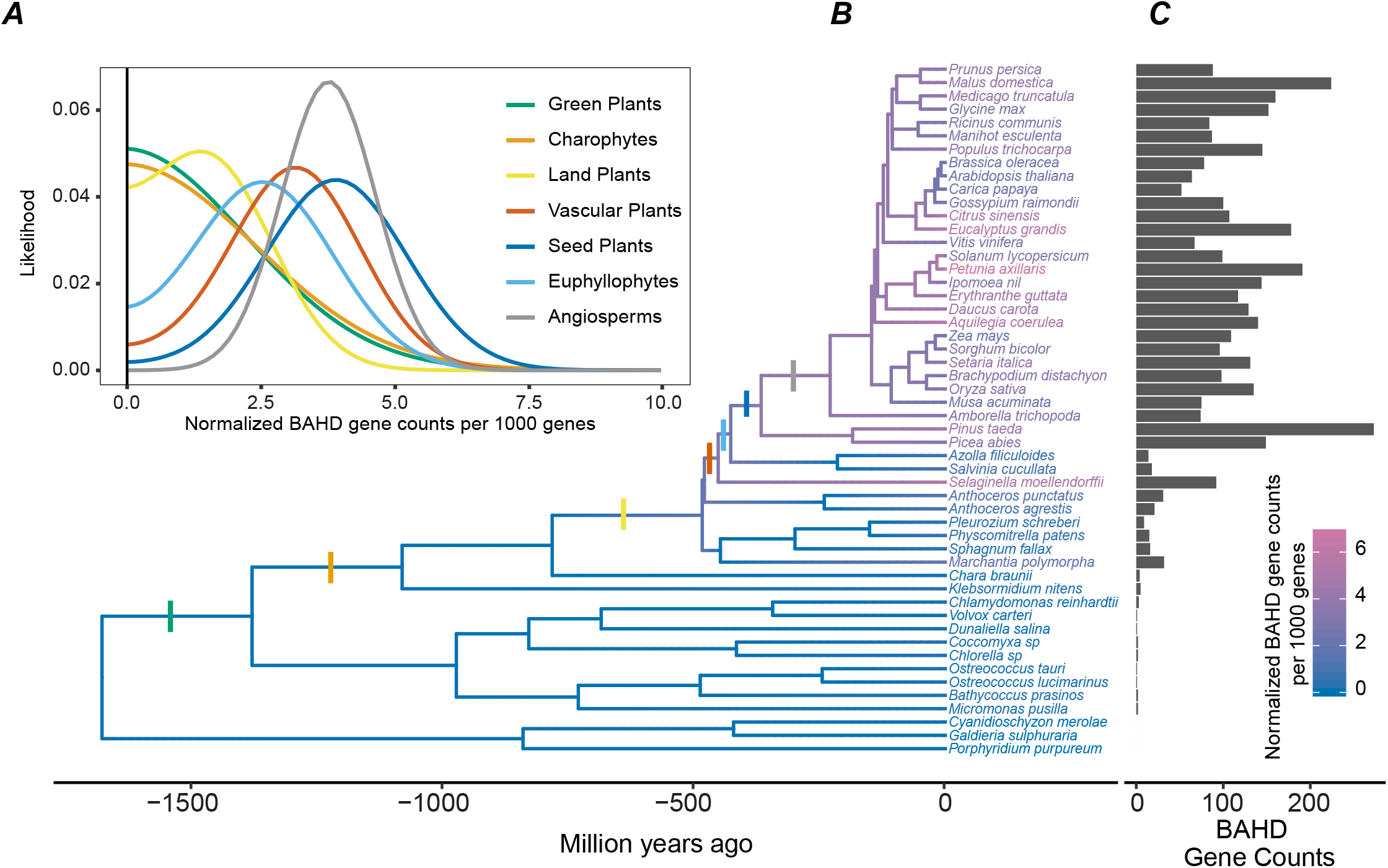
Major BAHD expansion occurred in land plants. **(A)** Likelihood of the number of BAHDs per 1000 genes in the genome of different phylogenetic groups, as per the bound Brownian motion model. The corresponding nodes are indicated in Fig. 3B. The bold, black line indicates the lower bound of the model which is set to 0 (no BAHD gene). **(B)** Species tree of the analyzed species with sequenced genomes. Color scale of the branches indicates the normalized BAHD count. **(C)** Bar graph showing BAHD count per analyzed genome.

Ancestral state function is typically obtained in two different ways – performing activity assays after ancestral sequence resurrection (Huang et al., 2016; Devamani et al., 2016) or inferring ancestral state based on analyses of activities of extant enzymes. However, BAHDs are fast evolving enzymes that maintain activity and structural folds despite undergoing large-scale sequence evolution and hence both approaches are not suitable for ancestral state reconstruction in this case (discussed in more detail in **Supp. Methods**). Thus, we used a different approach, where we first constructed 765 BAHD orthologous groups (OGs) across the 52 genomes, of which 132 OGs comprised >2 members and 89 OGs comprised members from >2 species (multi-species OGs). We then used sequence similarity to assign biochemically characterized enzymes to OGs, thus roughly predicting the OGs biochemical function at the substrate class utilization level. Depending on the breadth of conservation of a specific OG, we inferred the deepest internal node in the species tree likely housing the OG’s associated function, limited to dicots, monocots, and more internal nodes. More lineage-specific clusters (e.g. ASATs in Solanaceae) were defined manually in this study **(Fig. 4, Supp. Dataset 5A)**, but the phylogenetic resolution can be altered computationally as needed.

**Figure 4.**
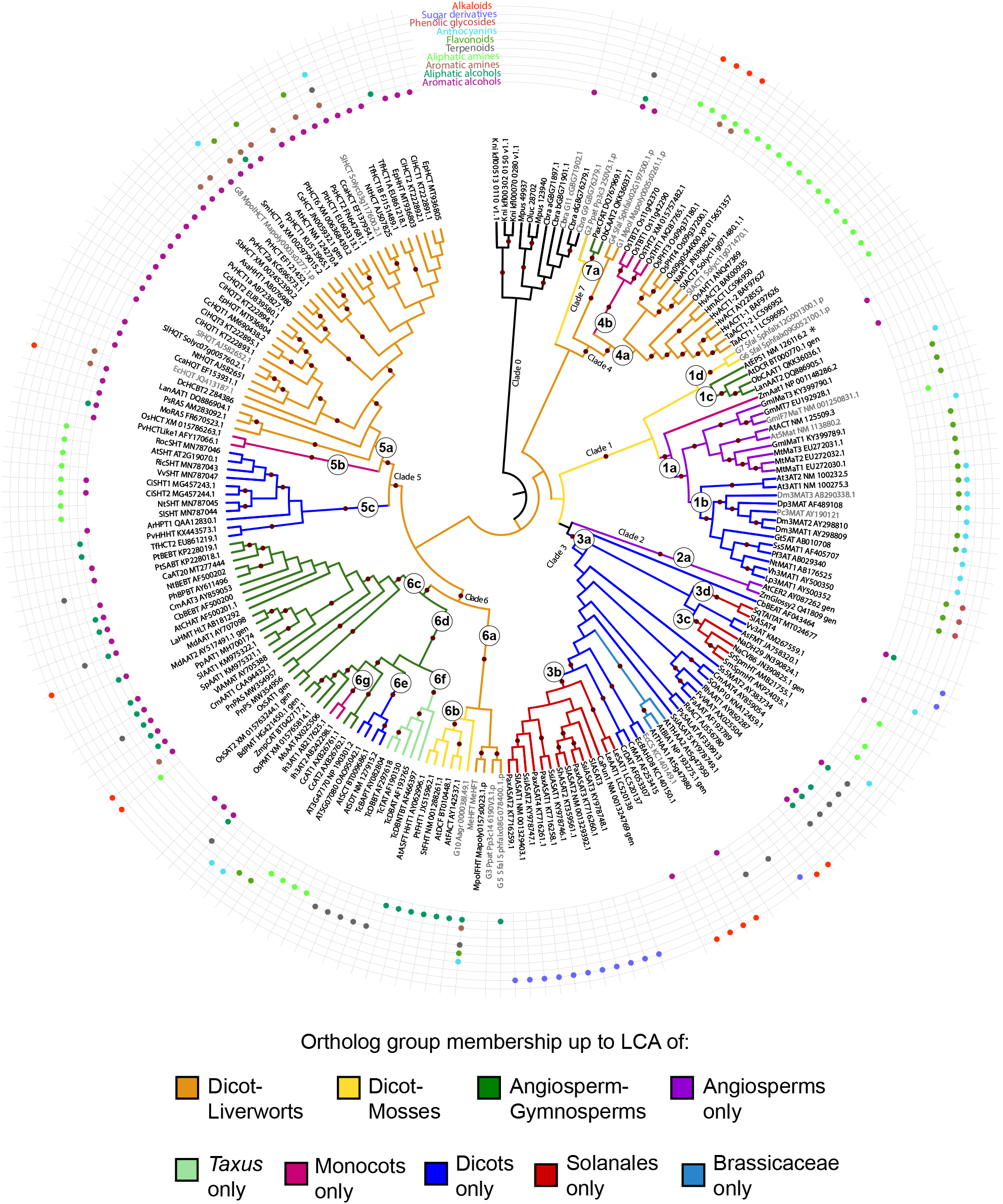
The eight clades of the BAHD family were established via duplications at multiple periods in the evolutionary history of plants. The tree was rooted using the algal enzyme clade (Clade 0). Maroon circles on branches refer to clades with bootstrap values > 70 (1000 bootstrap replicates). Clades were first defined based on deepest, high-confidence monophyletic clades. Clades 1-4 are same as D’Auria et al, 2006 definitions (Supp. Fig. 4), Clade V from that study is divided into Clades 5-6 here, and a new Clade 7 is loosely defined. Sub-clades were further defined within each clade based on the extent of conservation of OGs across the plant phylogeny (Supp. Dataset 5). These clade/sub-clade definitions can be further expanded as new, uncharacterized BAHDs are characterized in the future. Solid circles in the concentric circles around the tree represent activities characterized in this and previous studies, with “gen” referring to genetically characterized BAHDs whose substrates were not considered. Enzymes with names in gray are the ones assayed in this study. * AtEPS1 showes unusual reaction mechanisms for BAHDs and has an isochorismoyl-glutamate A pyruvoyl-glutamate lyase activity that produces salicylic acid.

This novel OG-based clade function annotation approach was validated separately using 28 previously characterized BAHDs **(Supp. Dataset 5C),** which revealed that 88-92% of these BAHDs could be correctly assigned a correct substrate-class based on the previously characterized activities in their OG group. The 164 biochemically characterized BAHDs **(Supp. Datasets 2 and 5)** were mapped to only 47 OGs, suggesting that ∼50% of the 89 multi-species OGs still have unidentified functions. Most internal node functional inferences were supported by multiple characterized BAHDs with the same function **(Supp. Dataset 5A)**, thus providing confidence in assigning the phylogenetic extent of most clades in the BAHD gene tree **(Fig. 4)**.

In addition to clade 0 that comprised uncharacterized algal enzymes, the previously characterized enzymes can be divided into additional seven clades with high bootstrap support, of which four (clades 1-4) are the same as defined previously (D’Auria, 2006). Clade V in D’Auria, 2006 was divided into two separate clades 5, 6 **(Fig. 4)**, and a new clade 7 consisting of “other” enzymes was defined. Three clades containing HCT/HQT enzymes (clade 5a), alcohol acyltransferases (clade 6a/b) and aliphatic amine acyltransferases (clade 4a) were the most widely conserved, with orthologs extending from angiosperms to mosses/liverworts **(Fig. 4)**, leading us to hypothesize that these functions existed prior to the origin of land plants. Based on branch lengths, Clade 5a is under the most purifying selection of all BAHDs, where most of its members are involved in phenylpropanoid pathway and lignin biosynthesis **(Supp. Fig. 4)**. This suggests that an unknown enzyme mapping to this clade using BLAST will likely share a similar function i.e., acylating substrates from the same substrate class. For example, the newly characterized enzyme SlHCT mapped to this clade/OG using BLAST and was able to perform aromatic alcohol acyltransferase function. Enzymes mapping to clades 1a/b (all but one anthocyanin/flavonoid acylating despite long branch lengths) and 6a/b (cuticular wax biosynthesis and slow-evolving) may also reliably perform anthocyanin/flavonoid and aliphatic alcohol acylation respectively. Other clades diverged rapidly at the sequence level, and thus it is difficult to predict their substrate class-level function based on simply their phylogenetic position **(Supp. Fig. 4)**.

### Testing the predictions of ancestral state of the family

The orthology-based analysis suggested that the ability to acylate aliphatic and aromatic alcohols and amines (orange clades) was already established in the ancestor of land plants **(Fig. 4)**. However, all of the *in vitro* characterized activities except PpHCT and MeHFT are from seed plants; thus, whether these activities exist in non-seed plants is not clear. Furthermore, the promiscuous activity of multiple HCT enzymes and MeHFT with anthocyanins/flavonoids **(Figs. 4, 2, Supp. Dataset 4)** also suggested the possibility of anthocyanin acylation being an ancestrally promiscuous activity that was later selected for in angiosperms. To test these predictions, we synthesized seven additional, previously uncharacterized enzymes from moss (*Physcomitrella patens, Sphagnum fallax*), hornwort (*Anthoceros agrestis*), and liverwort (*Marchantia polymorpha*) mapping to the above clades. Two additional clade 1 enzymes from *S. fallax* as well as two more divergent BAHDs from charophytic algae (*Chara braunii*) were also synthesized. Activities of these eleven enzymes were tested against representative substrates from aliphatic and aromatic alcohols and amines as well as anthocyanins **(Supp. Dataset 4)**.

Only 4/11 enzymes showed activities against the tested substrates. MpolHCT (clade 1) and the algal *C. braunii* enzyme were able to use aromatic alcohols. The *C. braunii* enzyme in clade 0 used quinate but not shikimate, leading us to label it as CbHQT based on its *in vitro* activity. Two additional enzymes from *A. agrestis* and *S. fallax* (clades 6a/b) were able to use decanol or 16-hydroxypalmitic acid as substrate, as predicted and in line with the MeHFT activity. Together with the phylogenetic analysis and presence of other independent activities, these results suggest aromatic and aliphatic alcohol acylation existed in BAHDs as primary activities prior to land plant origins.

The phylogenetic analysis strongly supported existence of amine acylation as ancestral, but the three enzymes we tested did not show amine acylation. One enzyme (MpolHCT) showed the anthocyanin acylation activity. Thus, four class A/B utilizing enzymes from non-seed plant lineages (MeHFT, SmHCT, MpolHCT) and three from seed plants (AtHCT, SlHCT, PsHCT2) showed the ability to acylate class D substrates. This activity also appears in at least six enzymes in other BAHD clades **(Fig. 4, Supp. Table 3)**. While independent evolution of anthocyanin/flavonoid acylation ability in each of these enzymes is a possibility, a simpler explanation is that the ancestral enzymes using class A/B substrates - especially HCT class of enzymes - may have had the capability to use anthocyanins promiscuously. This ancestral ability may have been selected in clade 1a/b, which almost exclusively uses anthocyanins/flavonoids. Comparisons of BAHD sequences in OGs could thus help identify specific residues that contributed to such activity shifts.

### Different residues in the catalytic site contributed to BAHD diversification

To identify residues important for activity shifts, we first compared the HCT/HQT- like enzymes (clade 5a/b) to amine-acylating enzymes (clade 4a), both clades being conserved across land plants. Thirty-five residues were present in >70% of the tested sequences in clade 5a/b but had completely switched to a different residue in >70% of the tested sequences in clade 4a **(Supp. Dataset 6)**. Of these, only two – AtHCT F303L and R356D **(Supp. Fig. 5A)** – were close to the active site, with R356 present in ∼90% of clade 5a/b but 0% of clade 4a, being replaced by Asp or Glu in a majority of clade 4a sequences. The R356D switch was previously shown to convert HCTs into using amine-containing substrates (Levsh et al., 2016; Chiang et al., 2018b). While this promiscuous activity exists in at least some *O*-acylating enzymes **(Supp. Fig. 3; Supp. Dataset 4),** the ubiquity of the R356D mutation in *N*-acylating enzymes suggests that this residue is critical for specialization towards positively charged substrates despite alternative binding sites in the protein contributing to increased promiscuity (Chiang et al., 2018a). For F303L, we found that while ∼95% of the HCT/HQT-like enzymes have the Phe residue, this is completely reversed, with ∼95% of the amine-acylating enzymes having a Leu residue. Although this residue was not experimentally tested, we hypothesize that this position also played an important role in diversification of the O-acylation activity.

Using OG comparisons, we also investigated sequence differences between HCT/HQT-like enzymes (clade 5a/b) and AnAT-like enzymes (clades 1a/b) using enrichment analysis. The OGs of clades 1a/b extend farthest back to angiosperms, which suggests that the fixation of the AnAT activity occurred only in this lineage **(Fig. 4)**. This spread corroborates with previous knowledge about evolution of the core anthocyanidin pathway in seed plants (Davies et al., 2020; Piatkowski et al., 2020), which is further extended via glycosylation, methylation and acylation. Anthocyanin acylation can improve the stability of the molecule to heat, higher pH, UV light – stabilizing flower color and providing more stable antioxidant properties in stress response – thus providing a possible evolutionary advantage to its fixation in flowering plants.

Motif enrichment analysis revealed two over-represented motifs in these clades (TFFDxxW: E-value=1.9e-248; YFGNC: E-value=4.5e-221**, Fig. 5A, Supp. Fig. 6)**. Single-residue analysis also found Trp36 and Cys320 (numbering following Dm3MAT3) in the two motifs that are conserved in AnATs in comparison to other biochemically characterized BAHDs (100% vs 0% and 95% vs 4%, respectively) **(Supp. Fig. 6; Supp. Fig. 5B,C)**. Both these residues were closer to the catalytic His than other identified residues in the crystallized structure of Dm3MAT3 in complex with malonyl-CoA (PDB: 2E1T) (Unno et al., 2007). Ancestral sequence reconstruction was performed to determine when these residues appeared in the BAHD phylogeny. Over 60% of residues in the ancestor of all AnAT-type enzymes could not be predicted confidently (posterior probability<0.5), owing to the rapid sequence evolution of BAHDs **(Supp. Dataset 7)**; however, both Trp36 and Cys320 were confidently placed in the ancestral node of clade 1a/b **(Fig. 5B**; posterior probability>0.95).

**Figure 5.**
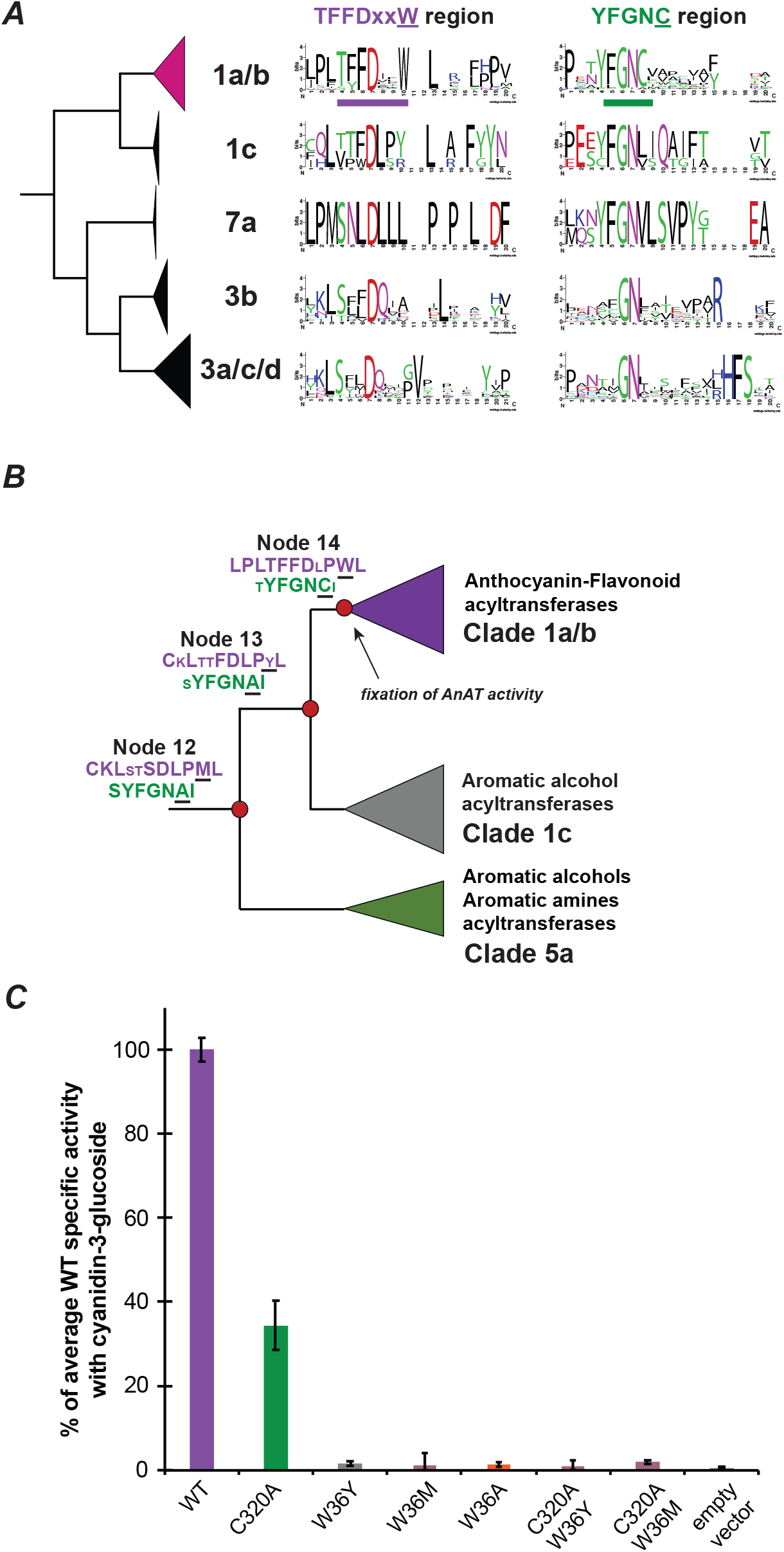
Trp36 and Cys320 are important residues for AnAT activity. **(A)** Conserved TFFDxxW and YFGNC motif region in different clades of the BAHD phylogeny. Clades are defined in Fig. 4. A 20 amino acid broad window is shown. **(B)** Ancestral sequence reconstruction of the conserved TFFDxxW (purple) and YFGNC (green) region. Large letters are residues with a posterior probability >80%. Smaller letters represent residues with a posterior probability <80%. Additional information can be found in **Supp. Dataset 7**. **(C)** Specific activity of Dm3MAT3 mutants. The average activi- ty of the wild-type (WT) is set to 100 %. The error bar represents the standard deviation of three replicates. The mutants W36Y and C320A-W36Y were assayed at a different time together with WT and empty vector control. The activity is normalized to the respective WT control that was run at the same timepoint. No malonyltransferase activity was seen in any enzyme with shikimate and decanol.

Given their near-ubiquity in clades 1a/b, we hypothesized that both these residues are critical for AnAT function. Emergence of Trp36 was most likely preceded by a Tyr (posterior probability<0.5) and Met (posterior probability>0.99), and a preceding Ala for Cys320 (posterior probability>0.99) in the prior ancestral nodes **(Fig. 5B; Supp. Dataset 7)**. To test if W36Y/M and C320A are functionally equivalent, we made single and double Dm3MAT3 mutants, as well as a non-aromatic W36A substitution to remove the spatial effect of the bulky ring. Comparing the specific activities of these mutants with anthocyanins as well as non-native substrates shikimate (aromatic alcohols, ancestral) and decanol (aliphatic alcohols, widely-distributed BAHD substrates) revealed that only the C320A mutation had significant, although substantially reduced, AnAT activity **(Fig. 5C)**. All other mutations led to almost or complete loss of AnAT function. These results supported our hypothesis that Trp36 is critical for anthocyanin acylation, however, Cys320 likely provides an accessory function for catalysis. Neither the wild type nor the mutants showed any promiscuous activities with non-anthocyanin substrates like shikimate or decanol.

These results also suggest that despite the large structural changes between the different clades that may have assisted with specialization of the AnAT activity, the acquisition of Trp36 and Cys320 was important for the angiosperm-restricted evolution of the activity. We next performed molecular dynamic (MD) simulations to determine the role of these residues in the AnAT activity. MD simulations were performed using the wild-type Dm3MAT3 enzyme and the W36A, C320A, and W36Y mutants.

### The role of Trp36 and Cys320 in anthocyanin malonyltransferase catalysis

The first step of the acyltransferase reaction involves proton abstraction from the cyanidin 3-*O*-glucoside (C3G) 6”-hydroxyl by the deprotonated, basic nitrogen of the His170 imidazole (**Fig. 6B**) (Unno et al., 2007). For successful intermolecular proton transfer to occur, the distance between these two atoms should be less than 4 Å to account for the longest possible hydrogen bond (Harris and Mildvan, 1999). MD simulations revealed the distance for C3G 6”-hydroxyl proton abstraction fell within a 4 Å threshold for WT Dm3MaT3 for the entire 1 µs of production runs. After maintaining a catalytically competent distance for the first 100 ns of simulation, the distance for proton abstraction in the C320A mutant exceeded the 4 Å threshold. The W36A/Y mutants never achieved a distance satisfactory for catalysis **(Fig. 6A)**.

**Figure 6.**
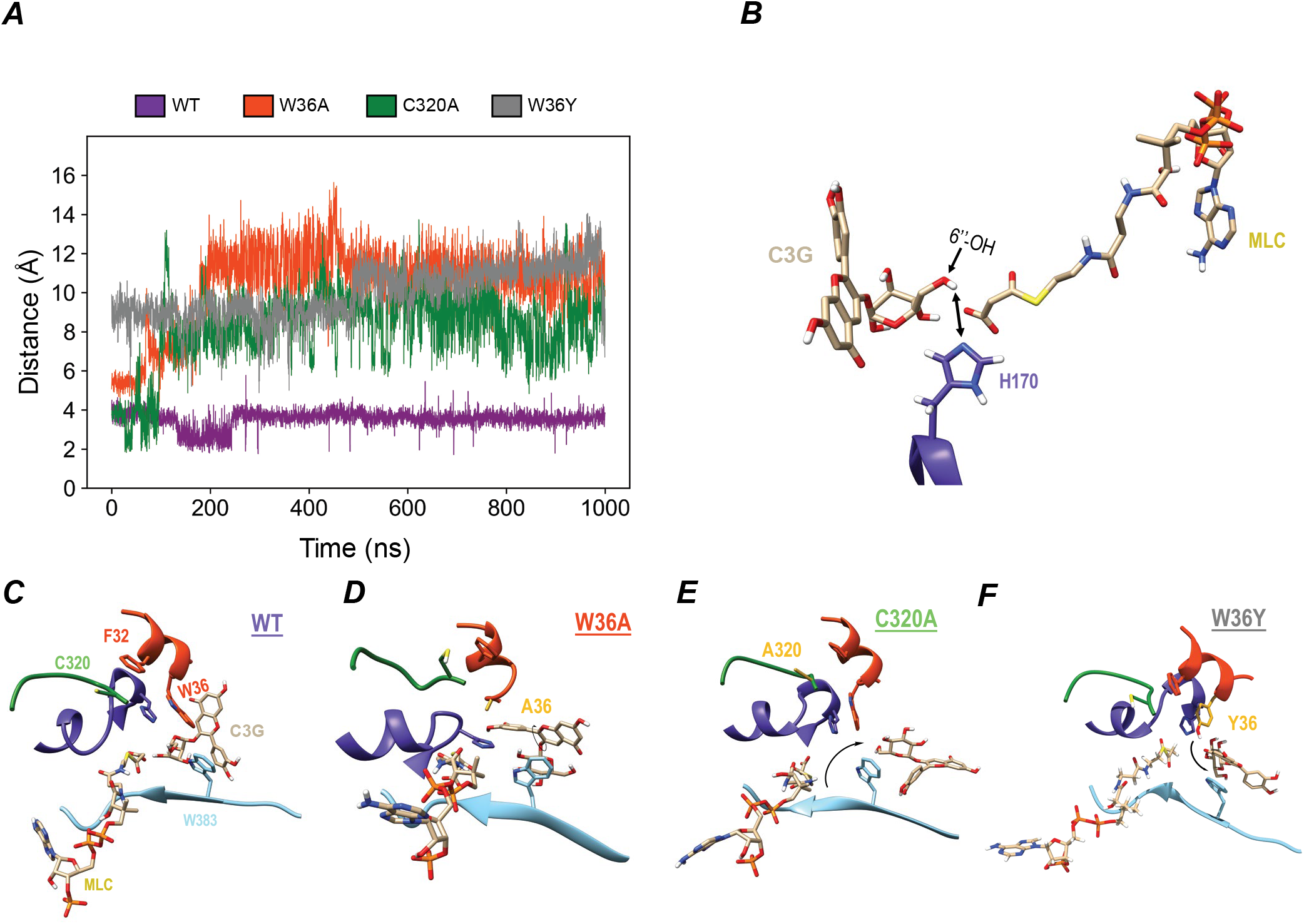
The residues Trp36 and Cys320 play important roles in maintaing the order of the activite site in Dm3MAT3. Distance calculations between the cyanidin 3-O glucosides 6”-OH hydrogen and the deprotonated, basic nitrogen of the His170 imidazole. **(A)** Distance calculation for WT (purple), W36A (orange-red), C320A (green), and W36Y (dark gray) over 1000 ns of simulation time. **(B)** Illustration of the distance measured during the simulation. Active site organization is altered upon replacement of Trp36 and Cys320. Active site organization in the (C) WT protein, **(D)** the W36A variant, **(E)** the C320A variant, and **(F)** the W36Y variant during catalysis. Substitutions leading to reduced enzyme activity are colored and labeled as goldenrod in the appropriate systems.

We next assessed how C3G is repositioned by each of the Dm3MaT3 variants **(Fig. 6C-F)**. In WT Dm3MaT3, the glucosyl moiety of C3G maintained a tight proximity to His170 within the active site, as it was flanked by bulky residues Trp36 and Trp383 **(Fig. 6C)**. The benzene diol moiety of C3G stacked with Trp383 and the chromenylium moiety stacked with Trp36. Having bulky residues in the proximal active site imposed a restriction on the movement of C3G, encouraging catalysis by keeping the acceptor ligand close to His170. Upon W36A substitution, the Trp bulk was absent, allowing C3G to sample other conformations within a more open active site while remaining bound to the enzyme. Trp383 initially stacked with aromatic portions of C3G in the WT, as seen in simulation. However, in W36A, the glucose moiety of C3G was flipped from its starting pose, where backbone hydrogens on the hexose ring demonstrate C-H···π interactions with Trp383, positioning the 6”-hydoxyl far from His170 **(Fig. 6D)**.

While the C320A mutant initially maintained a native active site organization due to the interactions between C3G and Trp36, C3G completely departed and was excluded from the active site by the end of the simulation **(Fig. 6A,E)**. Ala substitution altered the native C-H···π interactions between the backbone of residue 320 and the Phe32 benzene. In *apo* simulations, the Ala320 backbone hydrogen distance to the Phe32 π system was consistently less than 3 Å, but experienced fluctuations ranging Phe32 π between 2.5 and 6 Å in *holo* simulations **(Supp. Fig. 7A,B)**. For WT and W36A/Y variants, these distances were constant and unperturbed by ligand introduction, suggesting causes for reduced activity are different between the C320A and W36A/Y mutants **(Fig. 6D,F)**.

Fluctuations in the distance between the backbone alpha hydrogen of Ala320 and the Phe32 benzene were accompanied by a series of aromatic stacking rearrangements along the C320A variant TFFDxxW motif helix **(Supp. Fig. 7C,G)**. First, Phe32 became destabilized due to inconsistent backbone C-H···π interactions with Ala320, resulting in increased edge-to-face stacking between Phe31 and Phe32 **(Supp. Fig. 7A)**. To recover stability within the C320A variant’s TFFDxxW motif, Phe35 broke its edge-to-face stacking with Trp36 and began to stack with Phe31 and Phe32 **(Supp. Fig. 7D, Supp. Fig. 8)**. The W36A variant Phe35 never stacked with neither Phe31 nor Phe32 **(Supp. Fig. 9)**. Both WT and W36Y Dm3MaT3 also demonstrated virtually no Phe31-Phe35 nor Phe32-Phe35 stacking **(Supp. Figs. 10,11)**, suggesting these interactions were developed in response to instabilities from C320A substitution. Thus, Phe35-Trp36 stacking interactions were momentarily lost as Phe35 sampled new stacking interactions with Phe31 and Phe32 in the C320 variant **(Supp. Fig. 7D)**. With the network of stacking interactions between Phe35, Trp36, and His170 disrupted, C3G left the immediate vicinity of the C320A active site to bind elsewhere within the enzyme (**Supp. Fig. 7E,F**).

In contrast to the Ala substituted variants, wild-type Dm3MaT3 maintained an ordered active site which maintained better proximity to C3G and overall acyltransferase activity **(Fig. 6C)**. The stability and organization of the wild-type enzyme is underscored by consistent Phe31-Phe32 face-to-face stacking and Phe35-Trp36 edge-to-face stacking, neither of which were disrupted by Phe31-Phe35 or Phe32-Phe35 stacking at any point during the *holo* wild-type simulation **(Supp. Fig. 10)**.

Still, tight active site organization is not the sole requirement for AnAT activity, as the experimentally inactive W36Y substitution was able to maintain aromatic stacking like WT Dm3MaT3 (**Fig. 5C, Supp. Fig. 11**). W36Y mutation makes the Dm3MaT3 active site less hydrophobic, enabling the Tyr36 phenol to hydrogen bond with C3G in ways that Trp36 cannot. The WT Trp36 side chain sterically locks the C3G chromenylium core, restricting the glucosyl 6”-OH to come within 4 Å of His170 (**Supp. Fig. 12A**). Conversely, W36Y positions the C3G chromenylium core in an orientation orthogonal to that seen during WT simulations (**Supp. Fig. 12B**). Lacking the bulk of Trp, Tyr36 anchors the C3G chromenylium core only by hydrogen bonding to a deprotonated oxygen using its phenol group. In turn, the benzene diol and glucose moieties experience more conformational freedom to explore hydrophobic regions of the active site away from His170 (**Fig. 6F**).

Trp36 is thus important for keeping the C3G proximal to His170 because of its bulk, aromaticity, and hydrophobicity. The MD simulation results therefore suggested the W36A variant would have inferior reaction kinetics in comparison to the WT. In addition, the C320A mutant maintained C3G 6”-hydroxyl and the His170 imidazole within a distance of 4 Å for ∼10% of the total simulation time (about 100 ns) (**Fig. 6A**). Given the MD simulations depicted that C320A may position the acyl acceptor in a catalytically competent position, we were interested in how C320A kinetics differed from WT Dm3MaT3. Therefore, we tested the effects of Ala replacement of Trp36 and Cys320 using site specific mutagenesis and subsequent *in vitro* enzyme assays.

### The catalytic importance of W36 and C320 residues in anthocyanin acyltransferase activity

We heterologously expressed and purified mutagenized Dm3MAT3 variants from *E. coli*. These proteins showed similar folding behavior based on similar gel migration and retention time in size-exclusion chromatography compared to the wild-type protein **(Supp. Fig. 13A,B)**, suggesting that the mutation did not affect protein folding; however, both mutants substantially affected enzyme reaction kinetics when comparing specific activities (**Figs. 5C**). Comparing the pseudo-first-order reaction kinetics (**Fig. 7A,B**) of the wild-type Dm3MAT3 enzyme with the W36A mutant revealed that the mutation did not influence the acceptor K_m_ value (**Fig. 7C**) but drastically reduced turnover number (k_cat_) and catalytic rate (k_cat_/K_m_) by ∼97% and 97.5%, respectively (**Fig. 7D,E**). All three kinetic estimates were substantially reduced for the acyl donor malonyl-CoA (**Fig. 7C- E**).

**Figure 7.**
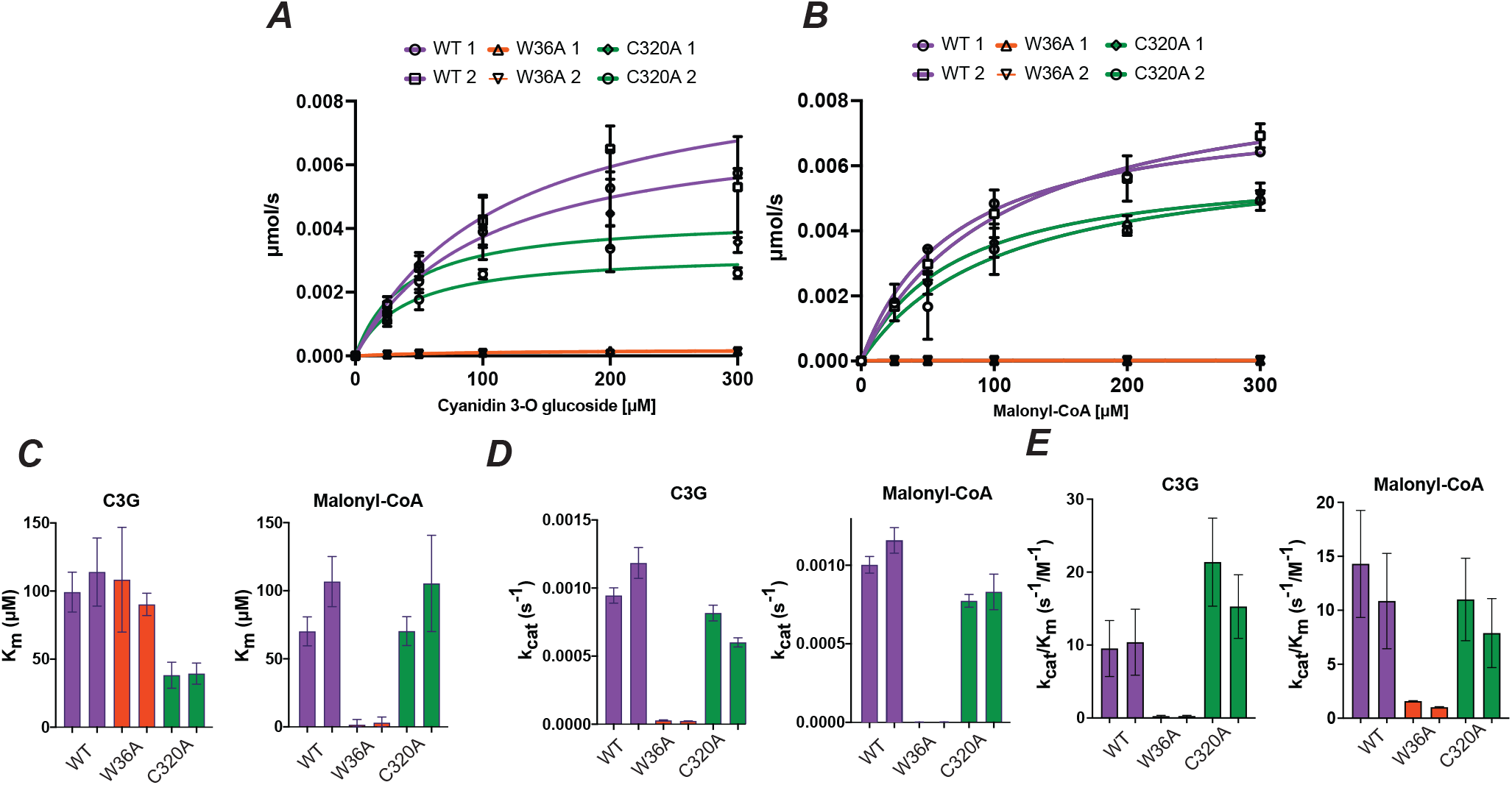
Alanine substitution of Trp32 and C320A severely affects enzyme catalysis. **(A)** Michaelis-Menten curve for Dm3MAT3 WT (WT), Dm3MAT3 W36A (W36A), and Dm3MAT3 C320A (C320A) in dependence of cyanidin 3-O glucoside (C3G). **(B)** Michaelis-Menten curve for the malonyl-CoA donor. **C)** Km values of WT, W36A and C320A. **(D)** k_cat_ values. **(E)** k_cat_/K_m_ values. For the data presented in panel (A) to (F), two independent expression replicates were performed and the kinetics were determined for each replicate individually using 3 technical replicates for each concentration. For each replicate the corresponding curve/bar is shown.

These changes in enzyme activity suggest that Trp36 is catalytically important for the AnAT activity in clades 1a/b. The relatively similar acceptor K_m_ between wild-type and W36A mutant suggested that the mutation did not affect acceptor substrate binding. Thus, while the MD simulations suggested a quick C3G departure from the active site, the molecule may still remain in the binding cavity due to interactions with other residues, and exit normally. The kinetic assays – specifically the reduced k_cat_ and k_cat_/K_m_ values for both acceptor and donor – support MD simulation predictions that catalysis would be drastically affected due to C3G departure from a catalytically competent distance with His170. The K_m_ of the W36A mutant towards malonyl-CoA was also substantially reduced. Considering AnAT-like acyltransferases operate through a two-substrate ordered bi-bi reaction mechanism with malonyl-CoA binding first (Suzuki et al., 2003; Unno et al., 2007), we postulate that the reduced catalysis in W36A results in the malonyl-CoA continually bound to the donor binding site, further resulting in a saturation of the available donor sites at lower substrate concentration.

In the case of the C320A mutant, the observed effects were less severe than for the W36A mutant (**Fig. 7A-E**), and the C320A still exhibited a k_cat_ ∼47-74% of the WT (**Fig. 7D**). Compared to the wild-type, the C320A mutant showed an improved K_m_ towards C3G (**Fig. 7B**). This lower K_m_ and the only slightly reduced k_cat_ resulted in a 50% improved catalytic rate than WT (**Fig. 7E**). The C320A mutant showed an unchanged K_m_ and a lower k_cat_ for malonyl-CoA. This resulted in a slightly less efficient enzyme with respect to the donor (**Fig. 7E**).

These results support the simulation prediction that Cys320 plays a role in optimizing the enzyme’s activity rather than catalysis. While MD simulations indicated that stability of the C3G bound form is reduced, this results in only a slightly decreased k_cat_. However, we postulate that the acceptor substrate remains in the substrate binding site without catalysis, reducing the K_m_ to a greater extent than reduction in k_cat_, thereby mathematically increasing the catalytic efficiency. Together, these results suggested that the presence of both Trp36 and Cys320 is necessary for optimal anthocyanin malonyltransferase activity in clades 1a/b, explaining their conservation in AnATs spread out over 140 million years of angiosperm evolution.

## DISCUSSION

Evolution of functional diversity in large enzyme families that contribute to the metabolic diversity in plants is still incompletely understood. In this study, we extensively characterized the BAHD family to determine the specific routes by which functional diversity arose in some clades of this family. We uncovered a broad prevalence and varied patterns of promiscuity among BAHDs. Nine out of fourteen BAHDs with activity detected in this study were substrate-promiscuous **(Fig. 2; Fig. 4; Supp. Dataset 4)**, and four (MeHFT, SlHCT, EcCS, MpolHCT) were class-promiscuous under the testing conditions **(Fig. 2A; Fig. 1)**. We only tested nine substrate types (including glucose/sucrose), and the specialized enzymes may still exhibit class- promiscuity with other untested substrates under different conditions. More BAHDs that cross class boundaries are known from previous studies **(Fig. 4; Supp. Dataset 2)**. Nonetheless, current evidence suggests that most BAHDs are substrate-promiscuous (e.g., shikimate, quinate for SlHQT, putrescine, agmatine for SlACT; **Supp. Dataset 4**) and are specialized enough to differentiate between very different chemical scaffolds. For individual enzymes, these adaptations for specialization may constrain them and their duplicates from traversing large distances in the phytochemical space. On the other hand, the class-promiscuity of enzymes such as SlHCT, AtHCT, SmHCT1, MeHFT, EcCS, MpolHCT, and alcohol acyltransferases **(Fig. 4)** may provide them/their duplicates the capability to traverse larger distances in the phytochemical space and “plug into” emerging metabolic pathways, forming a basis for metabolic diversification.

Significant presence of class-promiscuity can also confound evolutionary inferences and thus, functional annotation for enzyme family members based on sequence similarity. This issue has led to incorrect annotation of many uncharacterized BAHD family members in species’ genomes. For example, to the best of our knowledge in tomato, only 10 out of 100 BAHDs have an experimentally determined function (**Supp. Table 2**) (Niggeweg et al., 2004; Goulet et al., 2015; Fan et al., 2016) (including this study), and >80% of BAHDs have a generic domain annotation. Based on our phylogenetic analysis, we identify clades such as 5a (aromatic alcohol acyltransferases), 6a/b (aliphatic alcohol acyltransferases), 1a/b (anthocyanin/flavonoid acyltransferases) where substrate class-level functional annotation based on sequence is more likely to be correct. More characterized examples would be needed for other clades to offset the decrease in confidence due to the fast evolutionary rates of enzymes in those clades. The rise in utilization of machine learning algorithms for gene function prediction (Mahood et al., 2020) may enable better prediction of BAHD function using functionally relevant active site residues and knowledge of the substrate space accessible to the enzyme, despite high evolutionary rates in specific clades.

A central question of this study was to determine how the BAHD family came to use such diverse substrate classes. To answer this, we first inferred the ancestral state of the family prior to land plant evolution. We group the ancestral state prediction using three levels of confidence **(Supp. Table 3)**: **(1: High confidence)** Using previous enzyme information, new assays from non-seed plants, and phylogenetic range, we confirmed aliphatic and aromatic alcohol acylation (-OH acylation from class A/B, **Fig. 1**) as the primary ancestral activity prior to origin of the land plants. This inference is bolstered by previous predictions of aromatic alcohol acylation being ancestral, given its important role in monolignol biosynthesis in land plants (Weng and Chapple, 2010; Renault et al., 2019; Kriegshauser et al., 2021). **(2: Moderate confidence)** Five HCTs and one MeHFT – three of which are from non-seed plants – show promiscuous AnAT activities. Given the broad conservation of these clades and the high confidence in their primary activities being ancestral, we predict that the AnAT activity was also ancestrally existing as a promiscuous activity in HCT-like enzymes. Characterizing activities of more HCTs from non-seed plants will help bolster this inference. **(3: Weak confidence)** Based only on previous enzyme information, substrate structure similarity, and phylogenetic range, it is possible that the ancestral BAHDs were also able to acylate aliphatic and aromatic amines. Identifying amine acylation activities from liverworts, hornworts and algae can help increase confidence in this inference, but our studies failed to detect these activities. We also speculate that aliphatic alcohol-utilizing BAHDs may also use mono/sesqui/diterpenoids given the structural similarity of the substrates. Support for these inferences come from nine phylogenetically spread-out aliphatic alcohol acyltransferases that use terpenoids **(Fig. 4)**. However, more evidence from non-seed plants is needed to classify terpenoids acylation as ancestral (weak confidence). We did not explicitly test the latter hypothesis in our study.

The fixation of the AnAT activity in angiosperms is associated with fixation of the Trp36 and Cys320 residues in all clade 1a/b enzymes. We show that Trp36 is critical for the AnAT function, while the interaction of Cys320 with backbone groups helps in correctly positioning Trp36. Despite Cys320 being less proximal to the event of proton transfer than Trp36, it holds a structural role in shaping the active site. By stabilizing the TFFDxxW motif, Cys320 ensures tight organization at the site of reaction. Meanwhile, Trp36, with its bulk, aromaticity, and hydrophobicity, helps directly enforce a physicochemical environment where AnAT activity could emerge, as against Tyr-containing BAHDs in clade 1c. Together these residues affect acceptor binding; however, they are likely not sufficient to confer the AnAT activity. The median identity of AnAT enzymes in clade 1a/b is itself ∼50%, hence identifying all residues necessary to transform a class A/B utilizing enzyme to class D utilizing enzyme is challenging. Comparative sequence and structural analysis can help further identify regions in the protein that could be tested by mutagenesis.

Findings from this study also provide insights into the role of enzyme families in metabolic diversification in plants. BAHDs are fast-evolving enzymes; even in the well-conserved clade 5a, where all known enzymes are associated with substrates related to lignin biosynthesis (**Fig. 4**), the median percent identity is only ∼60%. The ability to use aromatic alcohols such as shikimate and quinate exists in BAHDs that are only 30-40% identical to the clade 5a BAHDs. In other words, these enzymes are robust in retaining their aromatic alcohol acyltransferase activity despite changes to ∼70% of their sequence. Similar behavior is observed for acylsugar acyltransferases (Moghe et al., 2017) and aliphatic alcohol acyltransferases **(Fig. 4)**. While being robust, the class-promiscuity of BAHDs and thus, the ability to specialize in one of the classes via duplication, would also make them evolvable. The paradox of a biological system being both robust and evolvable at the same time has previously been addressed in detail (Wagner, 2005; Bloom et al., 2006; Pigliucci, 2008; Wagner, 2008; Tokuriki and Tawfik, 2009; Payne and Wagner, 2019); these properties may enable enzyme families to become involved in newly emerging metabolic pathways or detoxify harmful metabolites without compromising their core activities. Presence of robust evolvable enzymes is likely an important feature of metabolic networks and needs to be studied in greater detail in the context of biochemical evolution and enzyme engineering (Irfan et al., 2021; Moghe and Kruse, 2018; Weng et al., 2021).

This study describes an integrative analysis of enzyme evolution, biochemistry and structure-function relationships that captures potentially emergent behaviors of enzyme families in the form of robustness and evolvability. While the findings reveal the extent of our knowledge on BAHD activities, there are 42 multi-species OGs not represented among characterized BAHDs, many of them showing broad conservation across angiosperms and vascular plants. These uncharacterized OGs may reveal novel and important BAHD activities. As new substrate classes are identified, it would be informative to test – using methods similar to those employed in this study – whether the ability to use those classes also exists across the BAHD family. Our results also identify patterns in duplication-divergence of BAHDs that can be explored in other large enzyme families to determine their involvement in metabolic diversification in plants.

## MATERIALS AND METHODS

### Creation of a database of biochemically characterized enzyme activities

Biochemically characterized BAHD enzymes were gathered through an extensive literature search. Only sequences belonging to the BAHD acyltransferase protein fold (PFAM domain: PF02458) were considered. To ensure a high level of confidence for *in vitro* activities and the resulting substrate-enzyme pairs, only enzymes that were subject to *in vitro* biochemical assays were considered. For each enzyme, all tested acceptor and donor substrates with their associated PubChem CID, the associated chemical structure, protein sequence as well as the species name from which the gene was isolated were compiled **(Supp. Dataset 1; Supp. Dataset 2)**.

### Generation of substrate similarity networks

Structures of known BAHD substrates were downloaded from the PubChem database using the PubChem ID as structure-data file (sdf) format. Substrates not found in the PubChem database were created using ChemDraw, exported in the MOL data format and manually brought into sdf format. To calculate substrate similarity based on the maximum common substructure (MCS), the R packages ChemmineR v3.34.1 and fmcsR v1.24.0 were used (Cao et al., 2008; Wang et al., 2013) with default values, except for the time threshold for the comparison of two molecules set to 12 s. The rationale for using MCS-Tanimoto is described in **Supp. Methods**. The similarity network was visualized using Cytoscape v3.8.0 (Shannon et al., 2003). Additional plant specific compounds were downloaded from the Chemical Entities of Biological Interest (ChEBI) database (Hastings et al., 2013) and only compounds were chosen that were also represented in the plant-centric KNApSAcK database (Afendi et al., 2012).

### Identification of BAHD acyltransferases and OG assignment

For identification of BAHD acyltransferases from the analyzed genomes, we used hmmsearch v3.1b2 (Potter et al., 2018) using the BAHD PFAM domain (PF02458) with all default parameters except *cut_ga* as the hit significance threshold. OGs were constructed using OrthoFinder v2.3.3 (Emms and Kelly, 2019) with default parameters except an inflation parameter of 1.5. After defining OGs between these species, we used blastp (Camacho et al., 2009) top hits to assign biochemically characterized enzymes to OGs. For each enzyme, its individual phylogenetic conservation was determined based on the phylogenetic spread of its assigned OG **(Supp. Dataset 5)**. An internal node in the species tree was assigned an activity if multiple characterized enzymes shared their conservation up to that node. While this method enables assignment of discrete states to specific ancestral nodes, there are no probability estimates associated with the predictions, which is a caveat of this approach.

### BAHD family size evolution

To determine the expansion dynamics of the BAHD gene family, we first created a time-calibrated species phylogeny (**Supp. Methods**) and used it to model the evolution of normalized BAHD gene counts. Counts were normalized against predicted total number of coding sequences in each genome with a methionine start residue that were longer than 100bp. If >30% of the gene models did not fit these criteria, these genomes (9 gymnosperm, 2 red algae) were eliminated from further analyses. We fitted normalized counts and our phylogeny using Evolutionary Brownian Motion (BM) and Bounded Brownian Motion (BBM) type models and performed model selection using the R package BBMV v2.1 (Boucher and Démery, 2016) with default parameters except minimum and maximum bounds for gene counts (see **Supp. Methods** for more details).

### Enzyme assays

In general, enzyme assays used for screening the substrate promiscuity of selected BAHD enzymes were performed in 50 µl reactions containing 50 mM sodium phosphate buffer (pH 7.2), supplemented with 100 µM acceptor substrate, and 300 µM donor substrate (substrate screening assays). Donor substrates p-coumaryl CoA and feruloyl CoA were biosynthesized separately as described in **Supp. Methods**. The assay was started by the addition of 1 µg purified enzyme. Mock controls using purified protein extracts from empty vector transformed *E. coli* were run to exclude *E. coli* background activity, for each of the used acceptor and donor combinations. For each enzyme, three replicates per substrate were performed and incubated for 1 hour. Reactions were stopped by adding 100 µl of a mixture of isopropanol, acetonitrile, and water (ratio of 2:2:1 + 0.1% (v/v) formic acid) containing 15 µM of the internal standard telmisartan. The reaction mix was centrifuged for 10 min at 21,000 g to remove precipitated proteins, transferred into LC vials, and stored at -20 °C until LC-MS analysis. Additional details regarding specific enzyme assay reactions and enzyme kinetic measurements can be found in **Supp. Methods**.

### LC-MS measurements

LC-MS analysis was performed on a ThermoScientific Dionex Ultimate 3000 HPLC equipped with an autosampler coupled to a ThermoScientific Q-Exactive HF Orbitrap mass spectrometer using solvent A (water + formic acid (0.1% v/v)) and solvent B (acetonitrile + formic acid (0.1%)) at a flow rate of 0.6 ml/min. Products of enzyme reactions were detected with specific PRM methods using their predicted parent ion mass in positive or negative ionization mode with an isolation window of 2 m/z **(Supp. Table 5)**. Additional details of chromatographic and mass spectrometric methods are described in **Supp. Table 6 and Supp. Methods**. LC-MS data was analyzed with the ThermoScientific Dionex Chromeleon 7 Chromatography Data System v7.2 software. Peaks were selected using their specific masses (**Supp. Table 5**) and default peak detection parameters.

### Phylogenetic analysis

To generate the tree shown in Fig. 4, a protein sequence alignment was generated using MAFFT v.7.453-with-extensions (Katoh et al., 2002) using following parameters: *--maxiterate 1000 --genafpair --thread 70*. The alignment was inspected manually to ensure proper alignment e.g., by inspecting that known motifs like the HxxxD and DFGWG motif are aligned properly. Afterwards, IQ-Tree v.1.6.10 (Nguyen et al., 2015) was used to infer a phylogenetic tree using following parameters: *-st AA -nt AUTO -ntmax 70 -b 1000 -m TEST* after model selection (LG+F+G4) using ModelFinder (Kalyaanamoorthy et al., 2017). Tree visualization was created using iTol v.5.6.2 (Letunic and Bork, 2019). Newick tree and display settings are available at: https://itol.embl.de/shared/gdm123

### Identification of enriched motifs in anthocyanin acylating enzyme

For identifying enriched motifs in a specific clade of anthocyanin acylating enzymes, we used MEME v.5.0.5 (Bailey et al., 2009) in discriminative mode using default parameters but the maximum number of motifs to find set to 5. The TFFDxxW and YFGNC sub-motifs were selected for further analysis due to their high degree of conservation and their proximity to the active site within the Dm3MAT3 protein. Clade-wise single residues were identified using custom Python scripts.

### Prediction of ancestral sequence of AnATs

Ancestral sequence reconstruction was performed using IQ-TREE v1.6.10 (Nguyen et al., 2015). Twenty randomly selected sequences from each of the three orthologous groups representing clades 1a-c and 5a (outgroup) were combined together with previously characterized BAHDs from these clades. All protein sequences were aligned using MAFFT v7.453 and provided as input to IQ-TREE, which was run with model selection and ancestral state reconstruction with 1000 standard bootstraps. The optimal tree was obtained using the JTT+I+G4 model. Per-site posterior probabilities of the ancestral state prediction were filtered using a threshold of 0.95.

### Molecular dynamics system preparation and simulation

Methods related to molecular simulations are condensed here and described in greater detail in the **Supp. Methods**. Briefly, the *holo* structure of Dm3MaT3 bound to malonyl-CoA (MLC), the acyl-donor, was retrieved from the Protein Data Bank (PDB: 2E1T) (Berman et al., 2000; Unno et al., 2007), treated with the PROPKA-plugin in VMD to verify residue charge states (Humphrey et al., 1996; Rostkowski et al., 2011), and then submitted to the Solution Builder Input Generator in CHARMM-GUI (Jo et al., 2008). The respective point mutations were then made during PDB structure manipulation. MD systems were constructed using the same pH conditions and ion concentrations as *in vitro* assays. MLC was taken directly from the 2E1T PDB file and was edited, along with Cyanidin-3-*O*-glucoside (C3G), in Maestro 2017-3 to reflect an accurate charge state (Schrödinger, LLC, 2020). Both ligands were parameterized in antechamber with the AM1-BCC charge model (Jakalian et al., 2002). Docking of C3G into each mutated Dm3MaT3 structure was performed using Autodock Vina v1.1.2, where more exact details concerning the docking protocol are provided in **Supp. Methods** (Trott and Olson, 2010).

Each *apo* and ligand-containing *holo* Dm3MaT3 system (WT, W36A/Y, C320A) was then reassembled as a single PDB in Packmol 18.169 (Martínez et al., 2009). Parameterization and periodic box conditions were then applied using tleap in Amber18 (Case et al., 2018). All ligands were parameterized with the GAFF2 (Wang et al., 2004), Dm3MaT3 variants with the AMBER-FB15 (Wang et al., 2017), and water with the TIP3PFB force fields (Wang et al., 2014b), respectively. Ions were parameterized using the 2008 parameter set developed by Joung and Cheatham (Joung and Cheatham, 2008). Hydrogen mass repartitioning (Hopkins et al., 2015) was then applied to the resulting files in ParmEd 3.2.0 (Case et al., 2018).

After initialization steps (minimization and heating), each of the eight systems underwent 1 µs of classical production runs (*holo* and *apo* versions of Dm3MaT3 WT, W36A/Y, and C320A). A Langevin thermostat was implemented for temperature maintenance (300 K) and a Monte Carlo barostat was implemented for pressure maintenance (1 bar) in the production runs (Loncharich et al., 1992; Åqvist et al., 2004). The SHAKE algorithm was applied to all stages of initialization except for minimization (Krautler et al., 2001), while the Particle Mesh Ewald method used for treating long-range electrostatics at a 10 Å cutoff (Darden et al., 1993). Python scripts were then written to calculate atomic distances and angles relevant to simulation analyses contained within the manuscript. Again, more details concerning system preparation, as well as postprocessing analyses, are described in the **Supp. Methods**.

## SUPPLEMENTARY MATERIAL

**Supplementary Figure 1:** Substrate classification and BAHD substrates represented using an alternative layout.

**Supplementary Figure 2:** BAHD substrates occupy a broad region in the phytochemical structural space

**Supplementary Figure 3:** Enzyme activities of selected enzyme representatives.

**Supplementary Figure 4:** Defining BAHD clades.

**Supplementary Figure 5:** Conserved residues in different clades of the analyzed BAHD acyltransferases

**Supplementary Figure 6**: Five topmost enriched motifs in anthocyanin/flavonoid- acylating (AnAT) BAHD acyltransferases (clade 1a/b).

**Supplementary Figure 7** Analysis of distance fluctuations in Dm3MAT3 variants.

**Supplementary Figure 8:** Stacking interactions for the C320A mutant.

**Supplementary Figure 9:** Stacking interactions for the W36A mutant.

**Supplementary Figure 10:** Stacking interactions for the WT Dm3MAT3.

**Supplementary Figure 11:** Stacking interactions for the W36Y mutant.

**Supplementary Figure 12:** Difference in cyanidin 3-O-glucoside binding to W/Y36.

**Supplementary Figure 13:** SDS-PAGE and Size Exclusion Chromatography of Dm3MAT3 variants.

**Supplementary Figure 14:** Substrate docking.

**Supplementary Table 1:** Substrate class definitions.

**Supplementary Table 2:** BAHD acyltransferases in the *Solanum lycopersicum* (tomato) genome.

**Supplementary Table 3.** Biochemical evidence for ancestral state of BAHDs.

**Supplementary Table 4:** Primer sequences.

**Supplementary Table 5**: Masses of detected products.

**Supplementary Table 6:** LC methods used for analyzing enzyme assay reactions.

**Supplementary methods:** Additional in-depth description of different analyses and experimental procedures.

**Supplementary Dataset 1:** Fasta file containing all analyzed BAHD sequences.

**Supplementary Dataset 2:** Description of compiled BAHD activities.

**Supplementary Dataset 3:** Identification of structural classes for compounds in the big network.

**Supplementary Dataset 4:** Measured enzyme activities

**Supplementary Dataset 5:** Mapping of biochemical characterized BAHDs.

**Supplementary Dataset 6:** Analysis of conserved residues between clades.

**Supplementary Dataset 7:** Ancestral sequence reconstruction.

**Supplementary Dataset 8:** Placing unresolved taxa and divergence time estimation.

**Supplementary Dataset 9:** Size exclusion chromatography (SEC).

## ACKNOWLEDGEMENTS

We thank Dr. John D’Auria, Dr. Toru Nakayama and Dr. Sangeeta Dhaubhadel for providing BAHD constructs, Dr. Bennett Fox and Dr. Frank Schroeder for use of and assistance with the LC-MS experiments. We thank Dr. Thomas Stegemann for support in LC-MS method establishment, Elizabeth Mahood and Dr. Nicholas Santantonio for support in establishing the substrate similarity pipeline, Dr. Hening Lin for helpful advice regarding our biochemical experiments, Dr. Elisabeth Kaltenegger for helpful comments on the manuscript, and Anna-Lena Sprick and Dr. Kai Fan for support in the laboratory. We thank Dr. Florian Boucher for assistance with interpreting BBMV R package output. This work was supported by the Deutsche Forschungsgesellschaft award # 411255989 (LK), Cornell University startup funds (GM), NSF award #DGE-1650441 (JMG and CS), NSF REU award #DBI-1850796 to the Boyce Thompson Institute/Georg Jander (JC), and the Blue Waters sustained-petascale computing project (NSF awards OCI-0725070 and ACI-1238993, the State of Illinois, and the National Geospatial-Intelligence Agency; AW and DS).

## AUTHOR CONTRIBUTIONS

LK and GM conceived and conceptualized the initial project idea. AW and DS designed and performed the molecular dynamics simulations; JMG, CS and GM designed and performed ancestral state reconstruction; JS and JJ designed and performed size exclusion chromatography experiments; JC and LK performed motif analysis; LK and M.I. performed enzyme assays; LK, AB, and MI performed LC-MS experiments. LK, MI and AB analyzed data LC-MS data, LK and GM performed and analyzed all other experiments. LK, AW, JMG, and GM wrote the manuscript and created figures. All authors reviewed the manuscript.

---

**Supplementary Figure 1:**
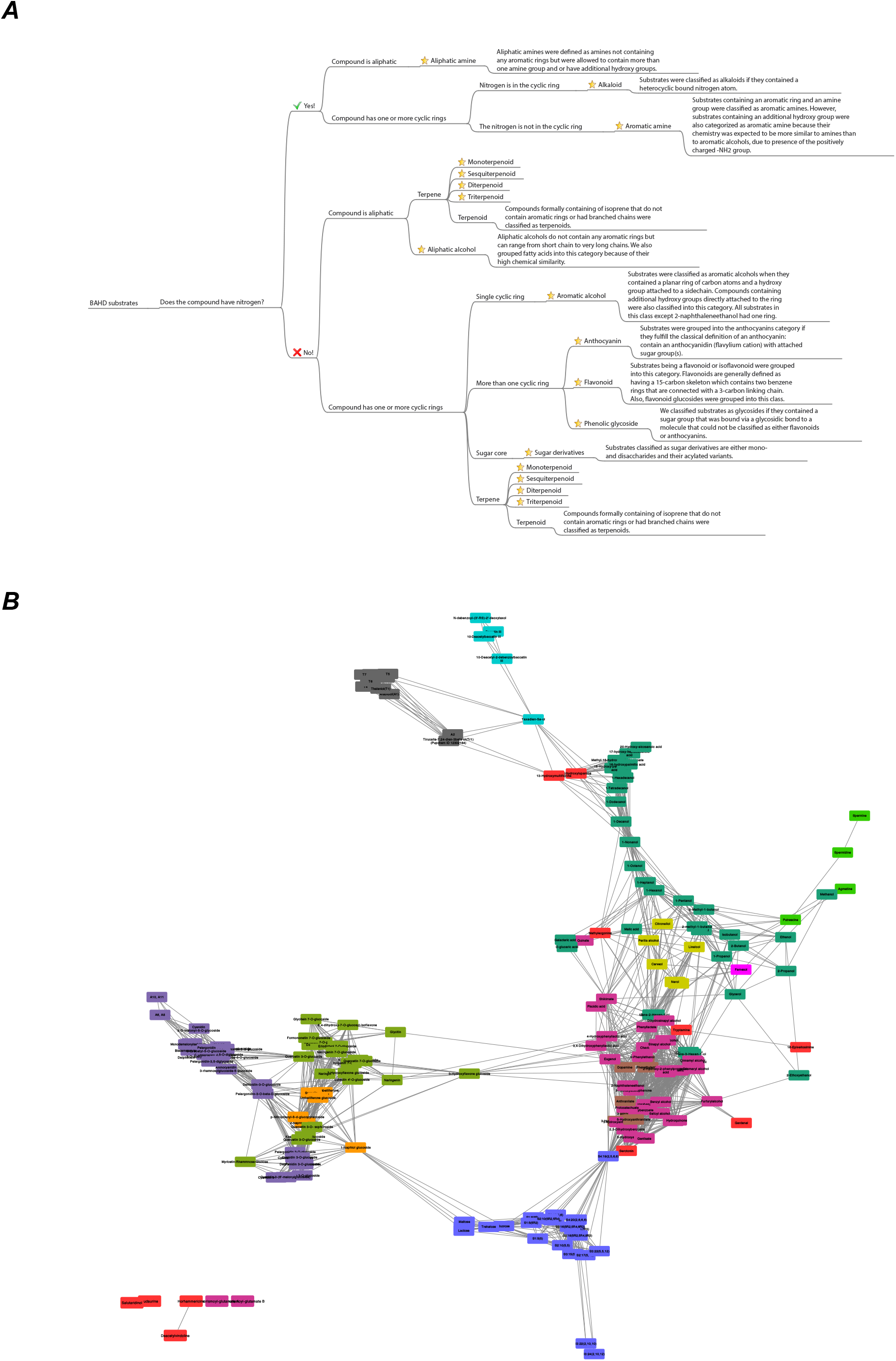
Substrate classification and BAHD substrates represented using an alternative layout. **(A)** Decision tree to group BAHD substrates into different substrate types. Further information about substrate class definitions can be found in **Supp. Table 1**. **(B)** The network was created in the same way and is colored the same way as Fig. 1. The “Edge-weighted Spring Embedded” Layout using MCS-Tanimoto coefficient was used to create an edge-weighted network.

**Supplementary Fig. 2:**
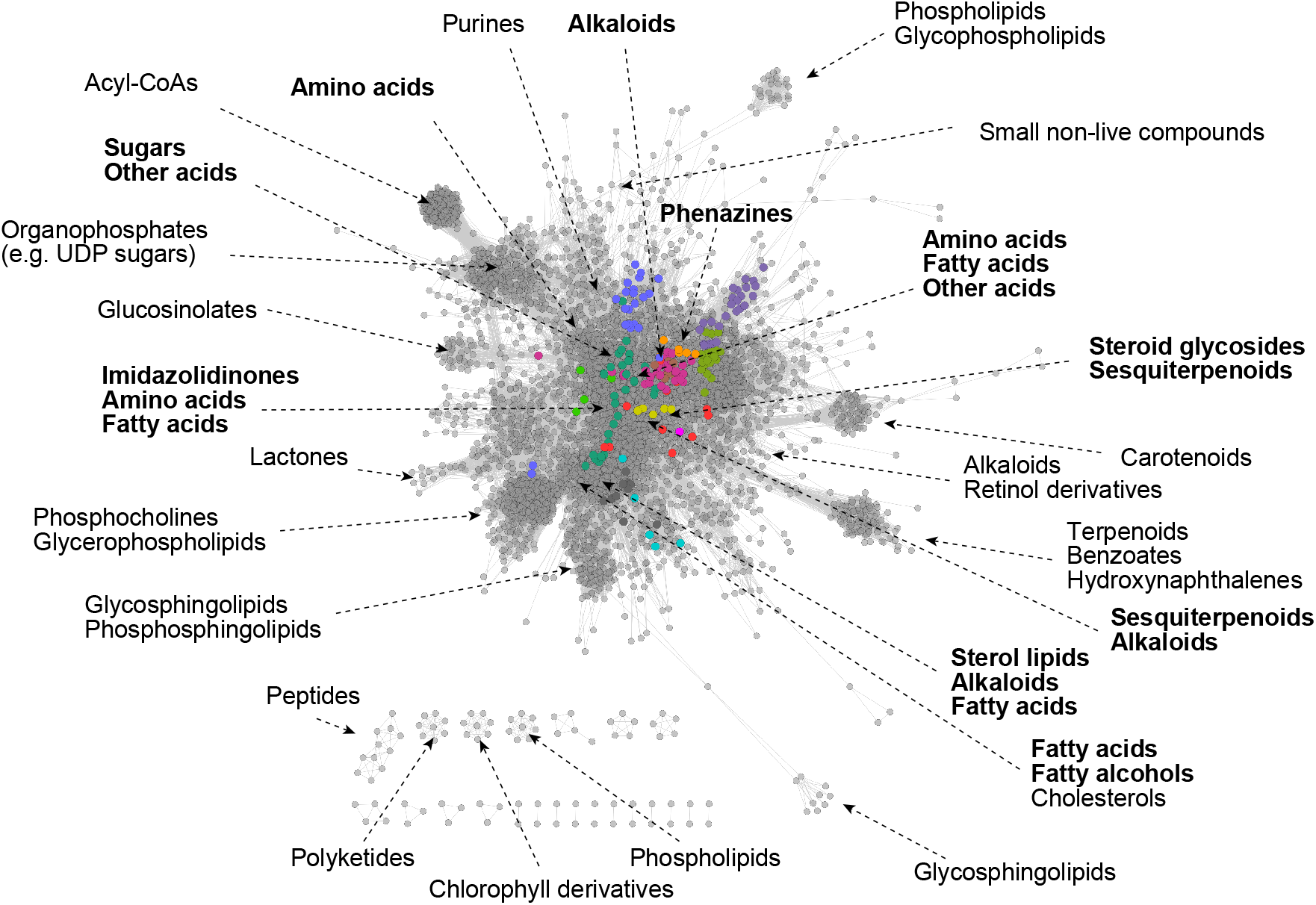
BAHD substrates occupy a broad region in the phytochemical structural space. Plant com- pounds with KNApSAcK ID were gathered from the ChEBI database. BAHD substrates are highlighted by colors corresponding to their substrate class. For each labeled cluster/region in the network, five compounds were chosen randomly and the compound class for each compound was determined based on the ChEBI Ontology (see **Supp. Dataset 3**). Representative classes are shown for each analyzed region. Names in bold are the classes closest to the known BAHD substrates.

**Supplementary Figure 3.**
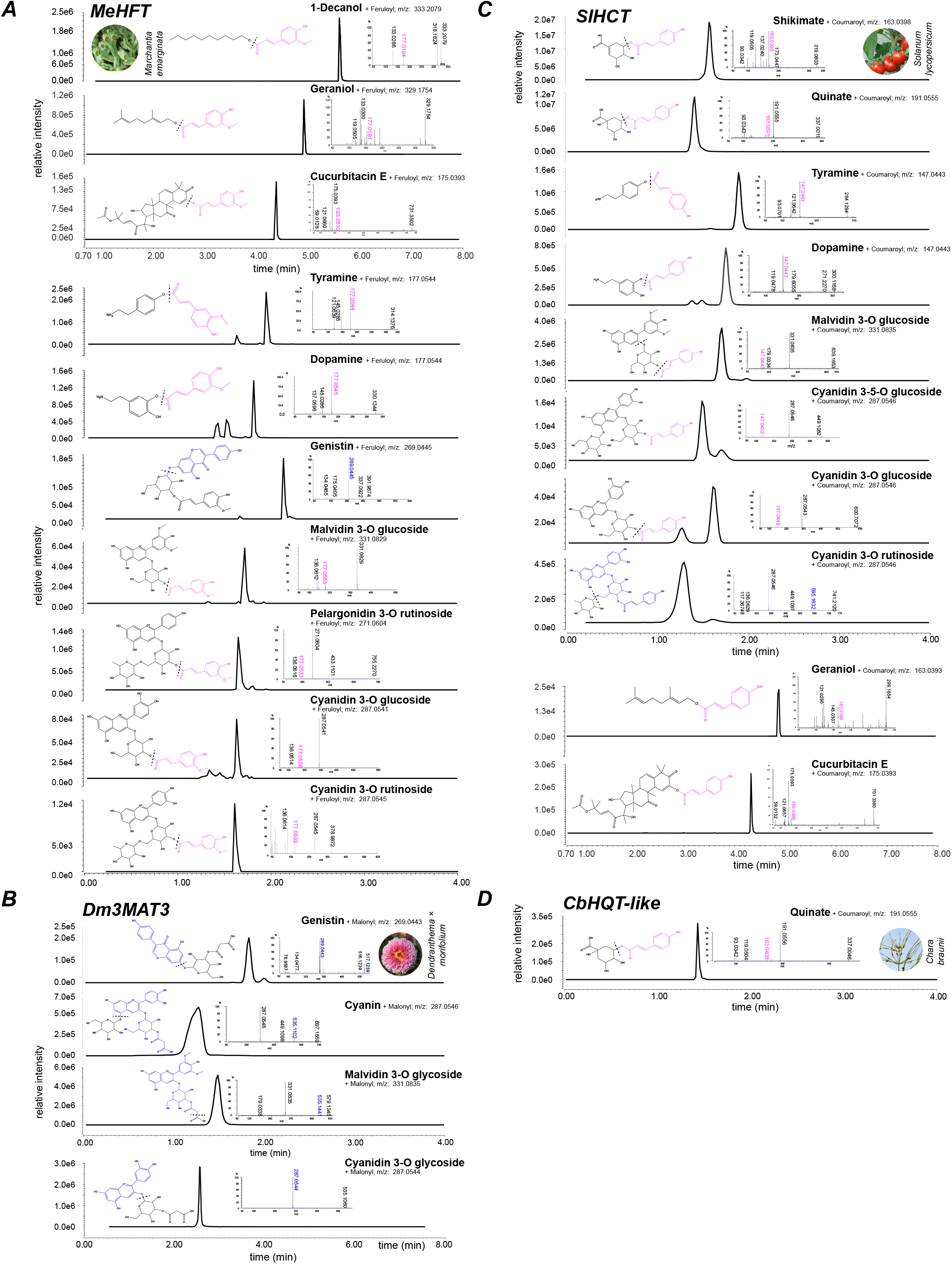
Enzyme activities of selected enzyme representatives. **(A-D)** extracted ion chromatograms of the quantifier ions of different enzymatic products of **(A)** MeHFT with its preferred donor feruloyl-CoA (**B)** Dm3MAT3 using malonyl-CoA **(C)** SlHCT using coumaroyl-CoA, and **(D)** CbHQT-like using quinate and coumaroyl-CoA. Each product was measured using product-specific PRM methods (see **Supp. Table 4**) and the most abundant fragment ion was used for quantification. This ion is noted in the upper right corner of each chromatogram. Structures represent the best-inference based on previously reported structures and the observed fragmentation patterns. Species photographs are from following sources: *Chara braunii* (Picture by Rob Palmer released under the CC BY-NC-SA license), *Marchantia emarginata* (Picture by Boon-Chuan Ho), *Dendranthema × morifolium* (Wikimedia Commons released into the public domain), and *Solanum lycopersicum* (no license attached).

**Supplementary Figure 4.**
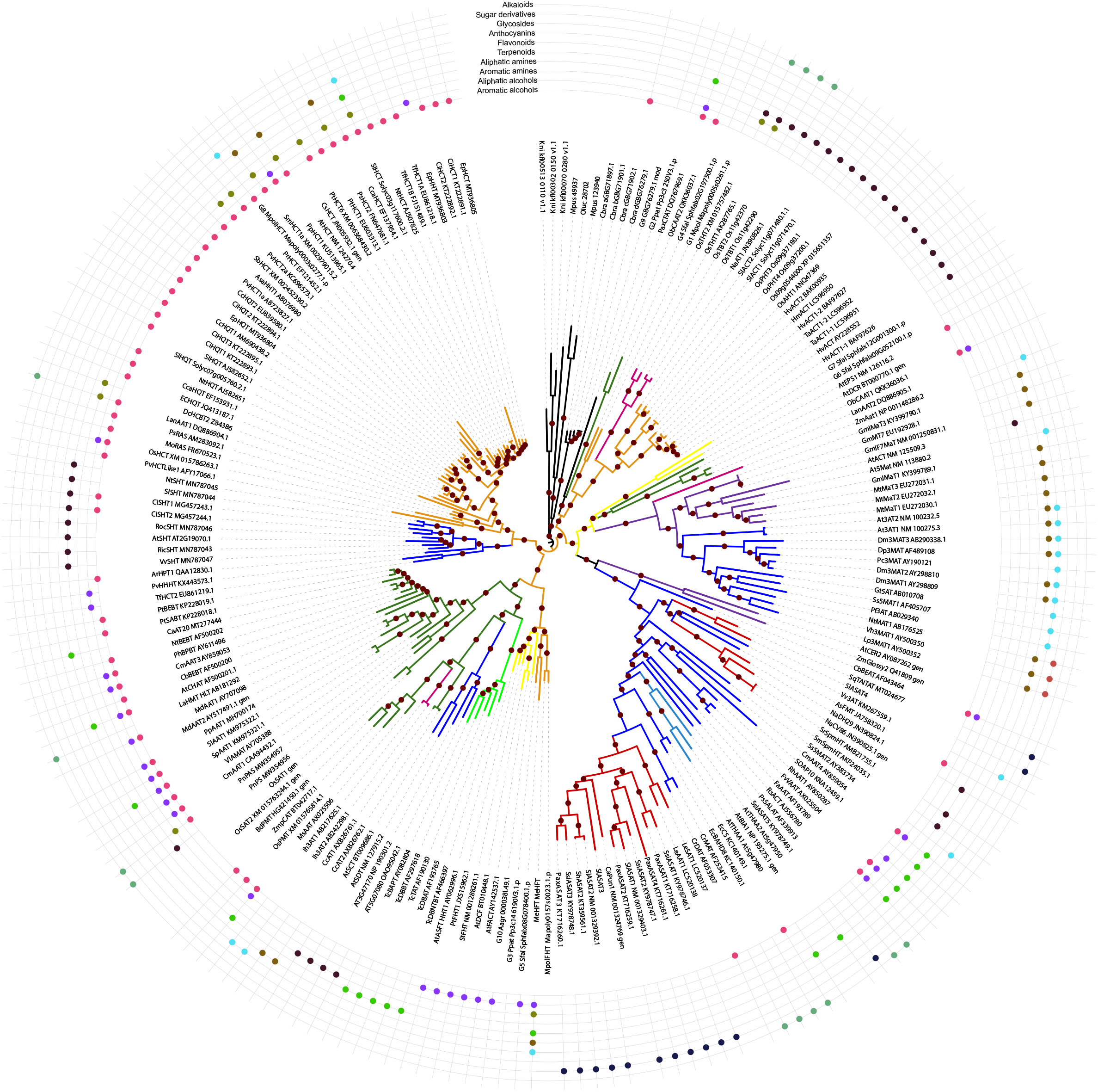
Defining BAHD clades. The tree was rooted using the algal enzyme clade (Clade 0a). Maroon circles on branches refer to clades with bootstrap values > 70 (1000 bootstrap replicates). Clades were defined based on deepest, high-confidence monophyletic clades. Clades 1-4 are same as D’Auria et al, 2006 definitions, while Clade V from that study is divided into Clades 5-6 here, based on the above criterion. Clade 7 is newly defined. Branch colors indicate the different clades. Branch lengths indicate number of amino acid substitutions per site.

**Supplementary Figure 5.**
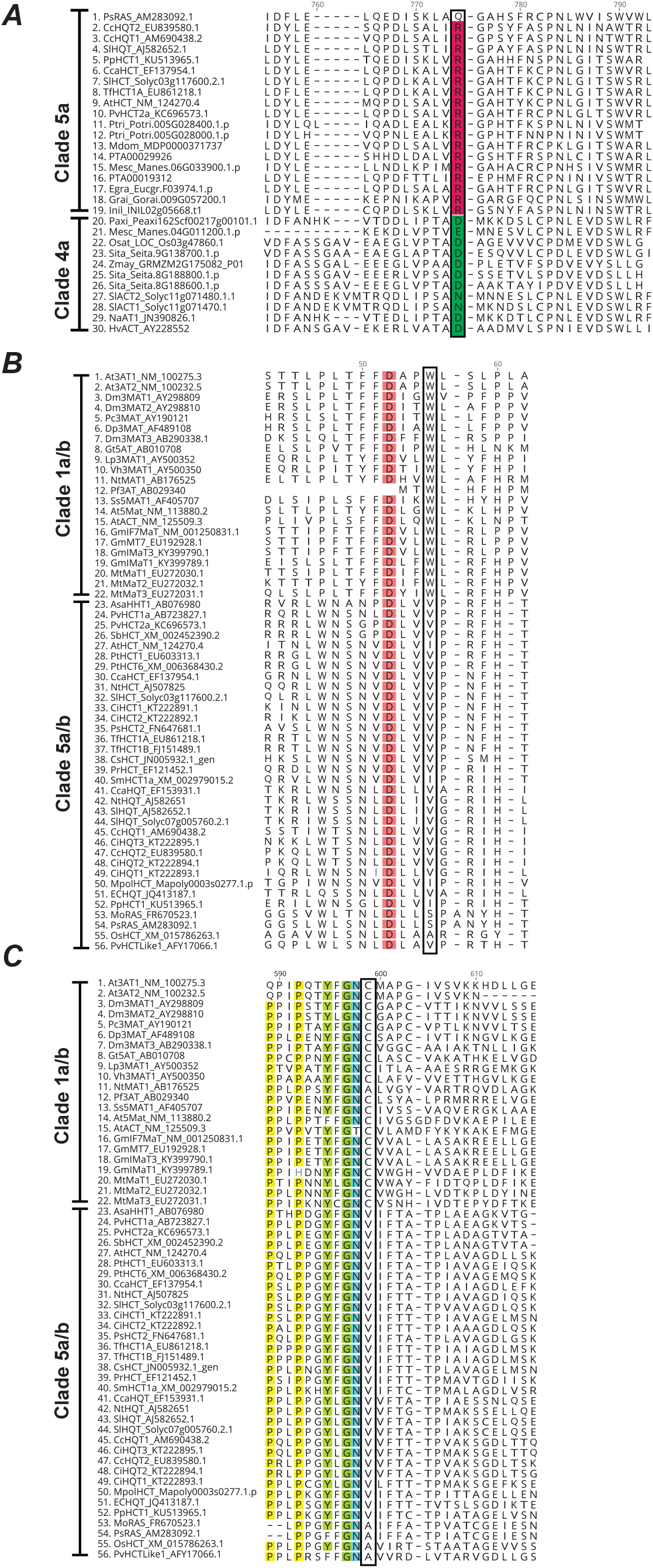
Conserved residues in different clades of the analyzed BAHD acyltransferases. **(A)** Alignment of clade 5a and clade 4a OG sequences. Highlighted in red is the Arg residue conserved in ∼90% of clade 5a BAHDs that predominantly use aromatic alcohols. In green, the corresponding Asp, Glu or Asn residues in amine acylating BAHDs are shown, of which Asp is present in ∼90% of clade 4a sequences. **(B)** Alignment of the TFFDxxW region of sequences belonging to clade 1a/b (anthocyanin/flavonoid) and 5a/b (HCT/HQT). **(C)** Alignment of the same sequences for the YFGNC region. Conserved (95%) residues are highlighted in colors. The positions of the highly conserved Trp and Cys residues are highlighted with a box.

**Supplementary Figure 6.**
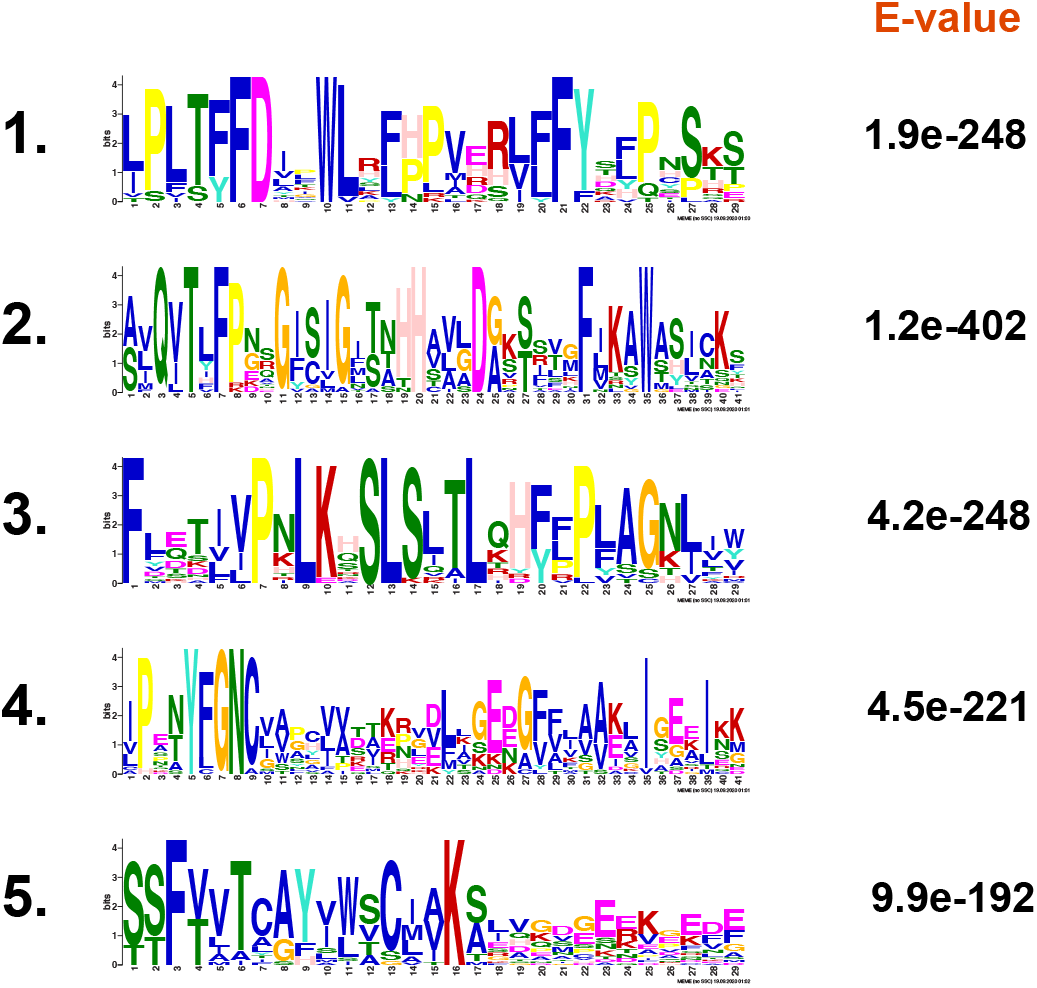
Five topmost enriched motifs in anthocyanin/flavonoid-acylating (AnAT) BAHD acyltransferases (clade 1a/b). For the analysis 22 AnAT BAHDs (clade 1a/b, Fig. 4) were compared against all other BAHDs from various clades.

**Supplementary Figure 7.**
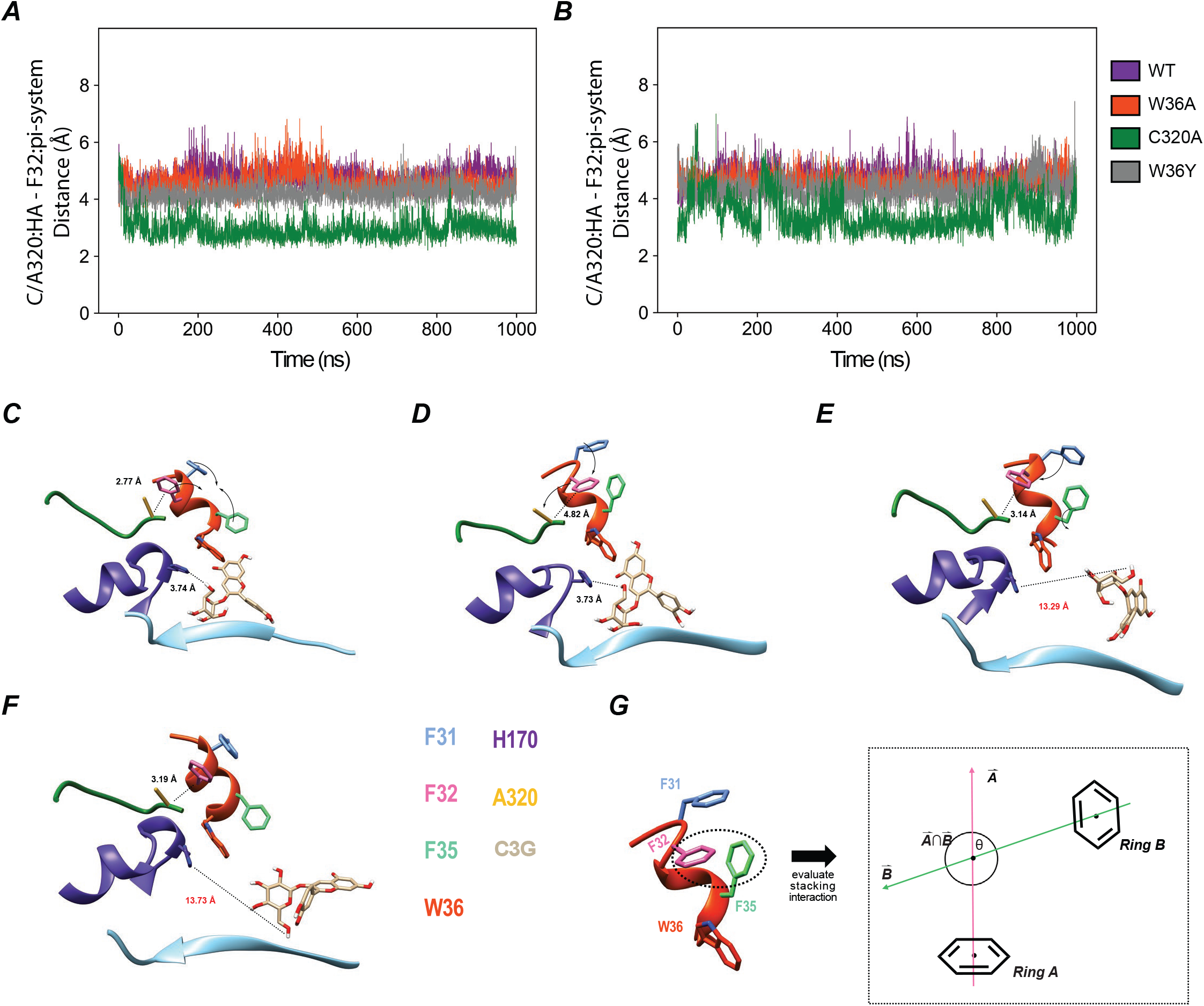
Analysis of distance fluctuations in Dm3MaT3 variants. **(A)** Distance calculations between backbone hydrogen of C/A320 and the center of mass of the aromatic ring of F32 for the apo simulation. WT is represented as purple, W36A is represented as orange-red, C320A is represented as green, and W36Y is represented as dark gray. **(B)** Distance calculations for the holo simulation. **(C)** Stacking rearrangements of phenylalanine aromatic sidechains within the TFFDxxW motif of C320A Dm3MaT3. Amino acid coloring: F31, cornflower blue; F32, hot pink; F35, spring green; W36, H170, A320, and C3G are colored as in previous images. Malonyl-CoA and W383 are hidden for simplicity. Distance fluctuations between A320 alpha hydrogen and F32 aromatic system cause aromatic stacking reorganization in the TFFDxxW motif. A catalytically competent distance between C3G-6”-OH and H170 is maintained. **(D)** C3G can still undergo reaction as F35 breaks its edge-to-face stacking with W36 to stabilize F31 and F32. (E) F31 and F32 have been stabilized. F32 moves closer to A320 while F35 returns to edge-to-face stack with W36, although C3G has distanced itself from H170. **(F)** Stacking interactions between residues in the active site and across the TFFDxxW motif have been reset, but now C3G is excluded from re-entering the active site due to steric and non-conventional bonding interactions with peripheral residues. **(G)** Schematic demonstrating θ calculations for discriminating between face-to-face and edge-to-face stacking interactions along the TFFDxxW motif. Figure and methodology as described in the Data Analysis subsection from the Main Text. Rings A and B represent the aromatic systems of two different residues. The normal vectors for each ring were then calculated so that the point of intersection between Rings A and B could be determined. Solving for the angle between the center of mass of Ring A, the center of mass of Ring B, and the intersection of the rings’ normal vectors, allowed for the supplementary angle, θ, to be determined. When θ ≤ 30° or 150° ≥ θ and the distance between the aromatic systems’ centers of mass is ≤ 4.4 Å, face-to-face stacking is occurring. When 60° ≤ θ ≤ 120° and the distance between the aromatic systems’ centers of mass is ≤ 5.5 Å, edge-to-face stacking is occurring. When 30° < θ < 60° or 120° < θ < 150° or the distance requirements for either form of stacking are unmet, no stacking interaction is occurring.

**Supplementary Figure 8.**
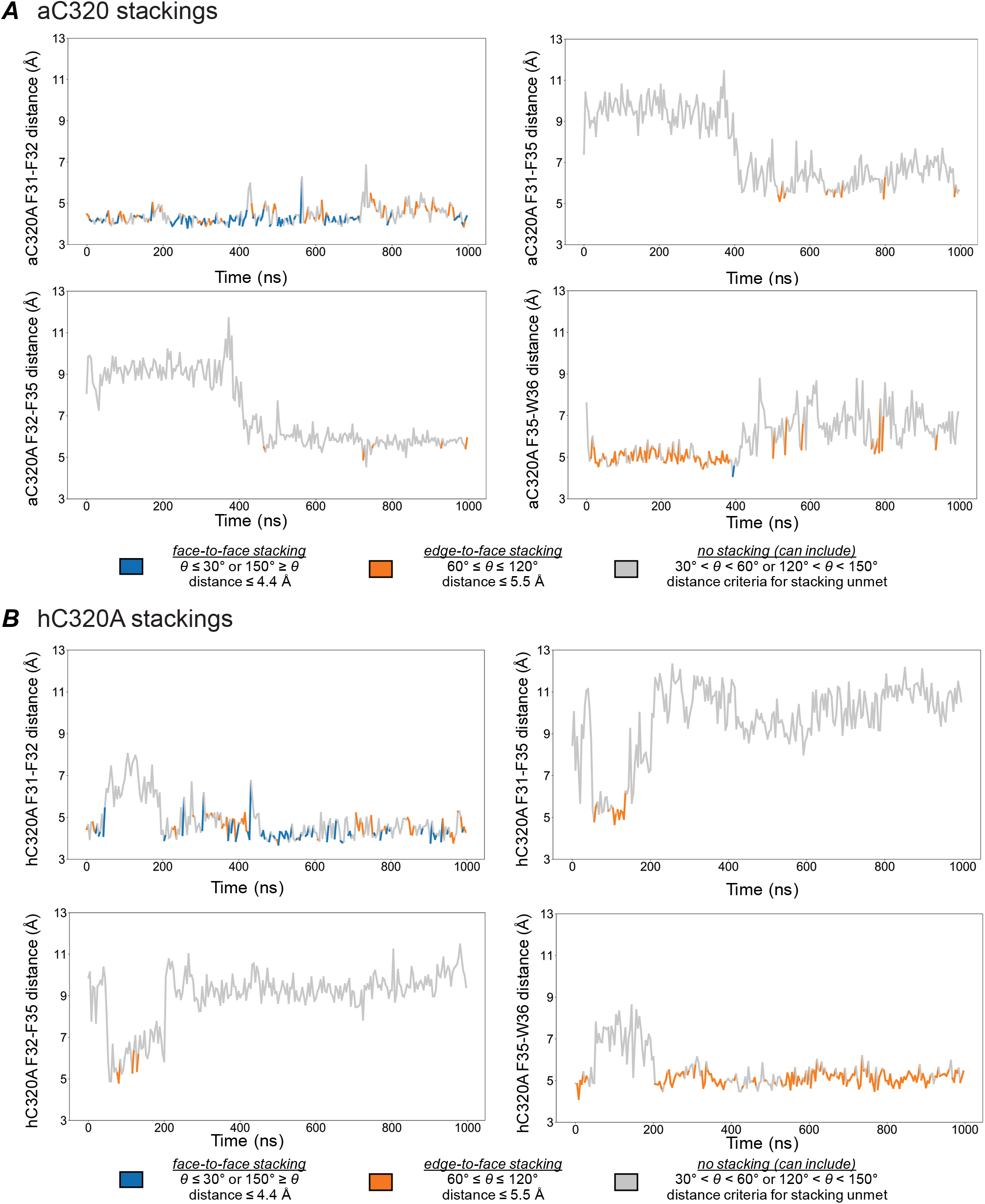
Stacking interactions for the C320A mutant. **(A)** Apo C320A (aC320A) stacking interactions categorized as face-to-face stacking, edge-to-face stacking, or no stacking interactions. **(B)** Holo C320A (hC320A) stacking interactions. Face-to-face stacking is represented as blue, edge-to-face stacking as orange, and no stacking as gray. Distance measurements were taken between the centers of mass of each residue listed on the y-axis of each panel.

**Supplementary Figure 9.**
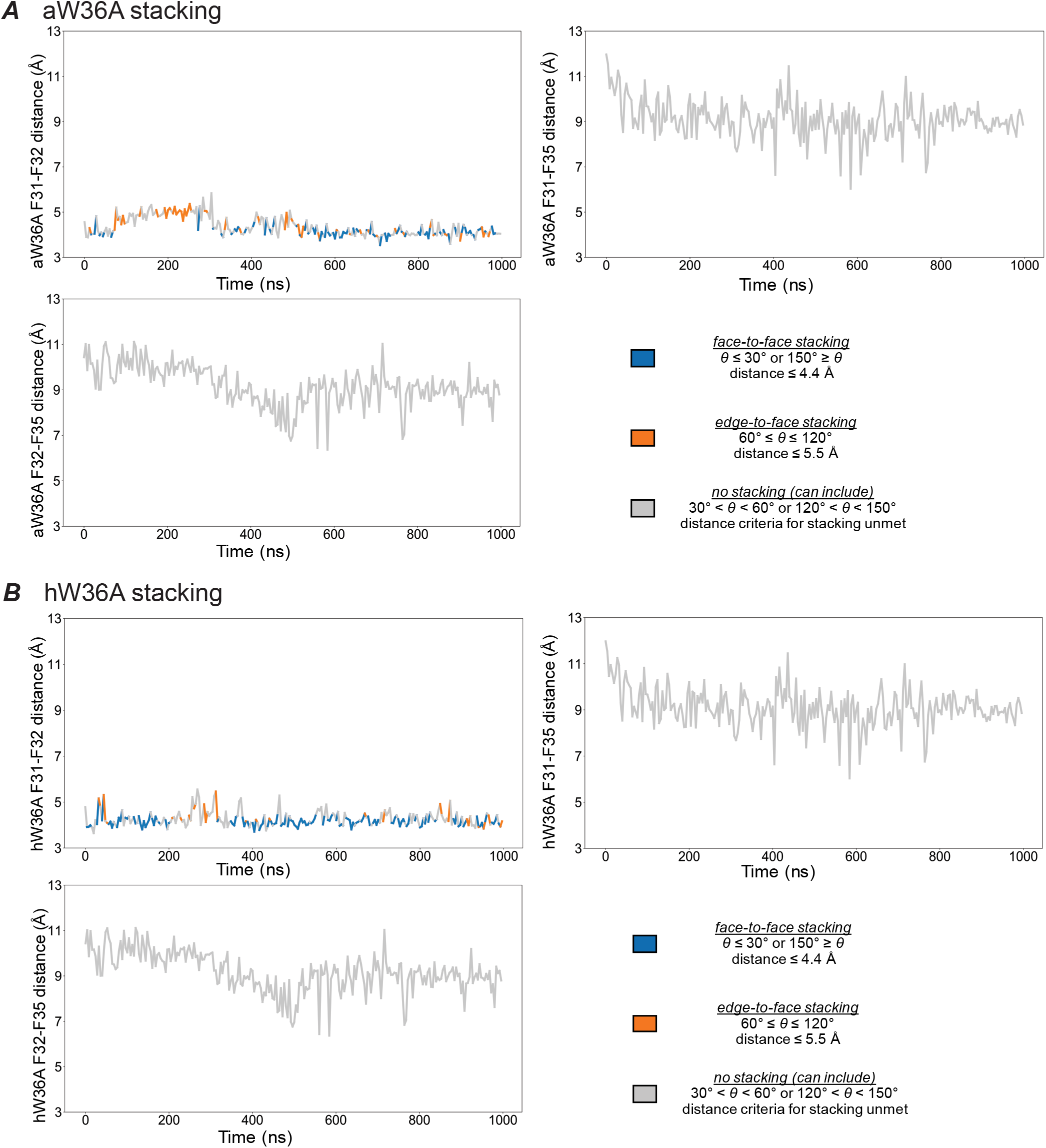
Stacking interactions for the W36A mutant. **(A)** Apo W36A (aW36A) stacking interactions categorized as face-to-face stacking, edge-to-face stacking, or no stacking interactions. **(B)** Holo W36A (hW36A) stacking interactions. Face-to-face stacking is represented as blue, edge-to-face stacking as orange, and no stacking as gray. Distance measurements were taken between the centers of mass of each residue listed on the y-axis of each panel. Only three panels are provided because W36A substitution prevents F35-W36 stacking from occurring.

**Supplementary Figure 10.**
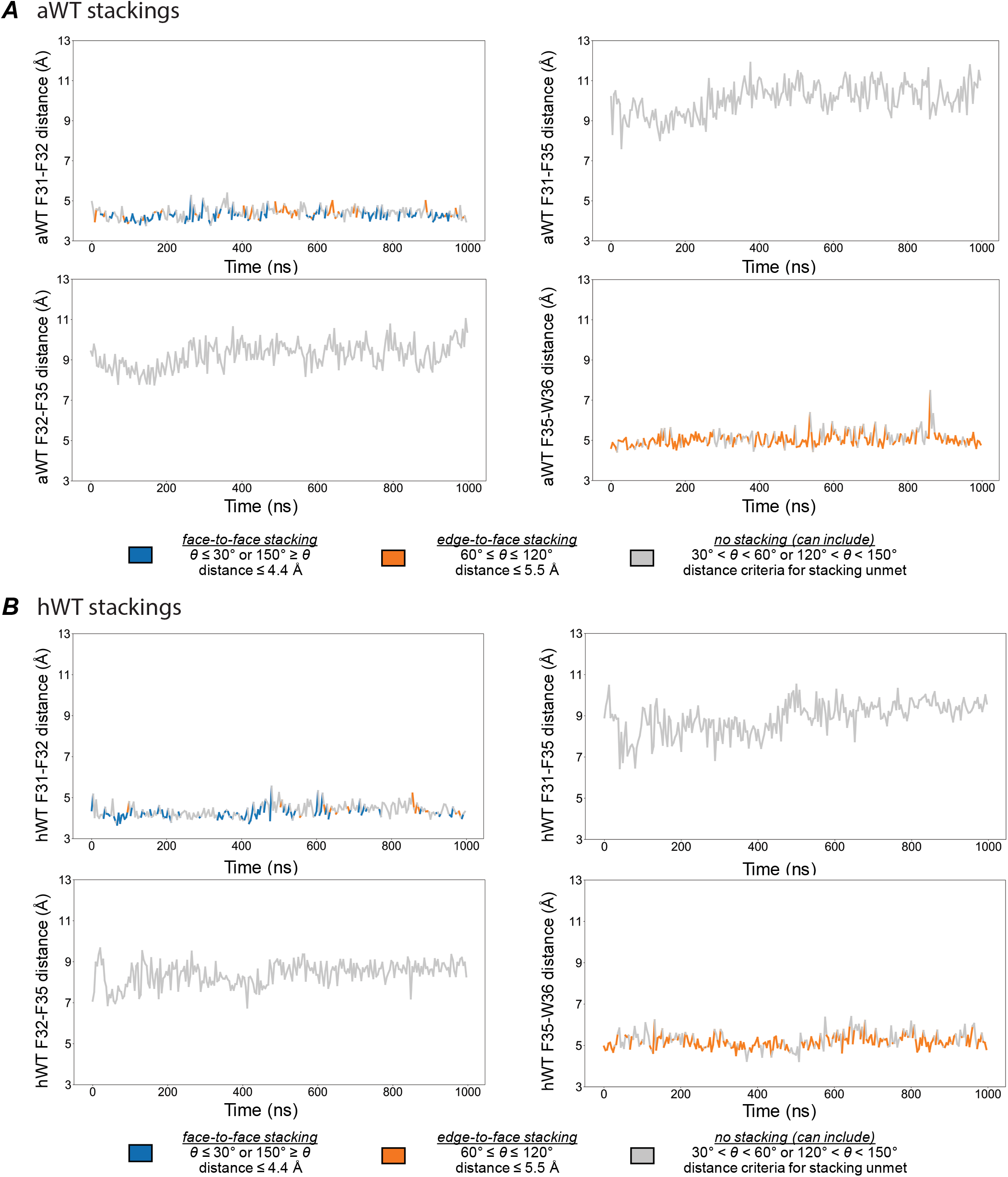
Stacking interactions for the WT Dm3MAT3. **(A)**Apo WT (aWT) stacking interactions categorized as face-to-face stacking, edge-to-face stacking, or no stacking interactions. **(B)** Holo WT (hWT) stacking interactions. Face-to-face stacking is represented as blue, edge-to-face stacking as orange, and no stacking as gray. Distance measurements were taken between the centers of mass of each residue listed on the y-axis of each panel.

**Supplementary Figure 11.**
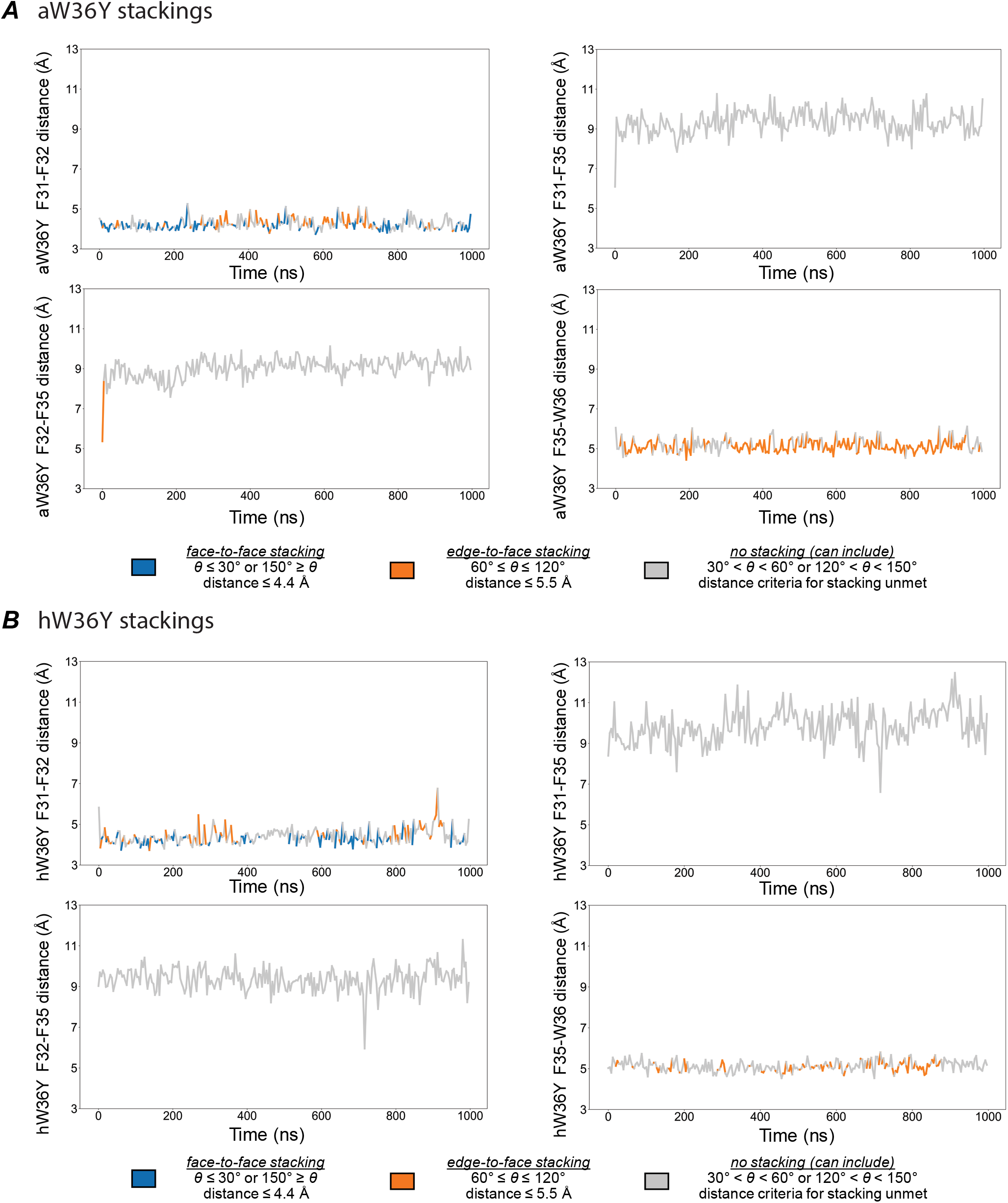
Stacking interactions for the W36Y mutant. **(A)** Apo W36Y (aW36Y) stacking interactions categorized as face-to-face stacking, edge-to-face stacking, or no stacking interactions. **(B)** Holo W36A (hW36A) stacking interactions. Face-to-face stacking is represented as blue, edge-to-face stacking as orange, and no stacking as gray. Distance measurements were taken between the centers of mass of each residue listed on the y-axis of each panel.

**Supplementary Figure 12.**
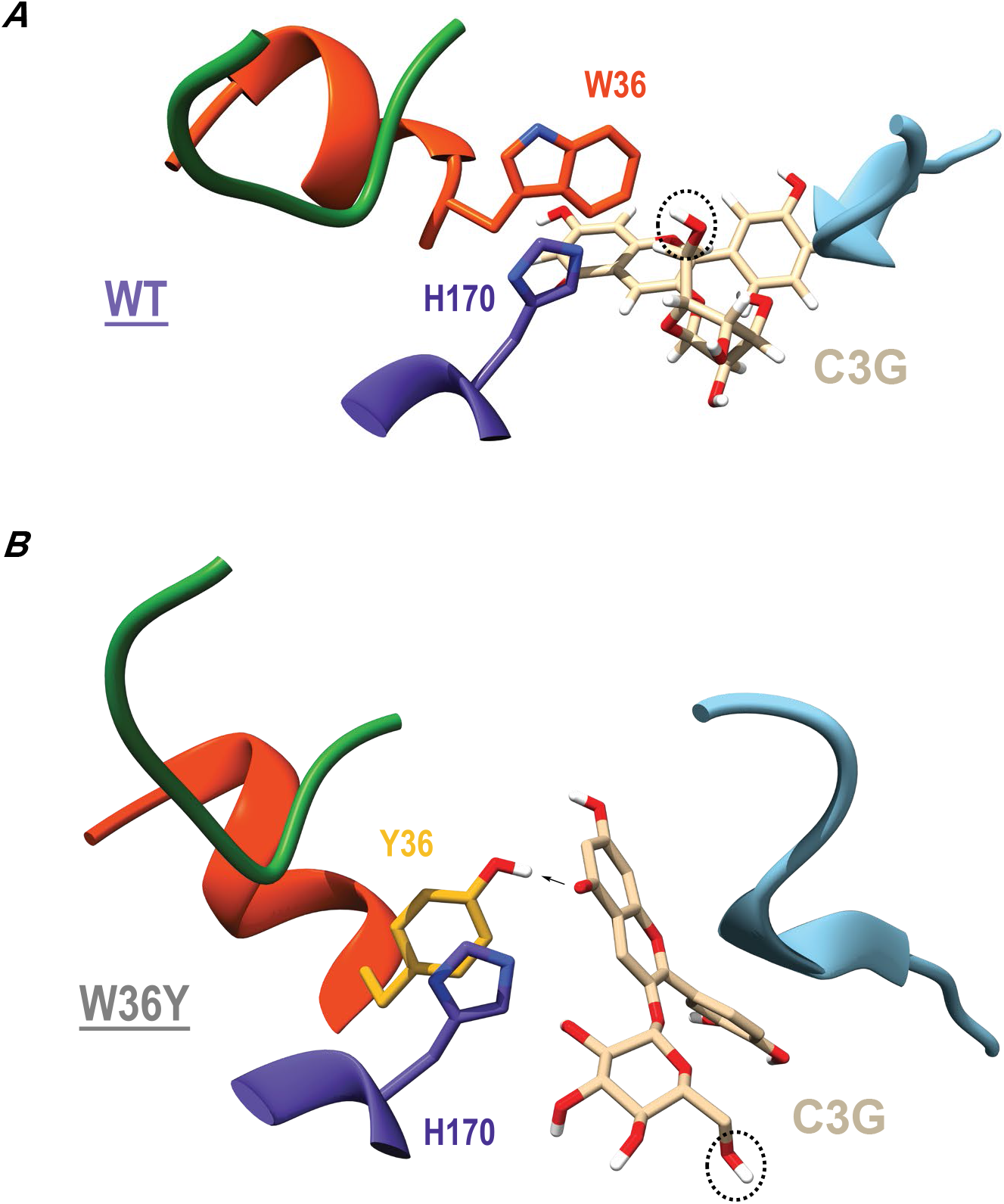
Difference in cyanidin 3-O-glucoside (C3G) binding to W/Y36. **(A)** W36 in WT Dm3MaT3 possesses the bulk to bisect the C3G scaffold core, trapping the chromenylium moiety in a perpendicular orientation based on the W36 bulk and hydrophobicity. This interaction reduces the conformational freedom of C3G, resulting in the reactive 6”-OH being within a catalytically competent distance throughout the 1 µs simulation (highlighted by dashed circle). **(B)** Y36 in the W36Y mutant lacks the bulk to trap the C3G chromenylium core. Instead, a hydrogen bond between the Y36 phenol and a deprotonated oxygen on the C3G chromenylium core serves as the dominant interaction, creating a C3G orientation orthogonal to that seen in WT simulations. Accordingly, the reactive 6”-OH experiences more conformational freedom and prefers a position ∼10 Å away from H170.

**Supplementary Figure 13.**
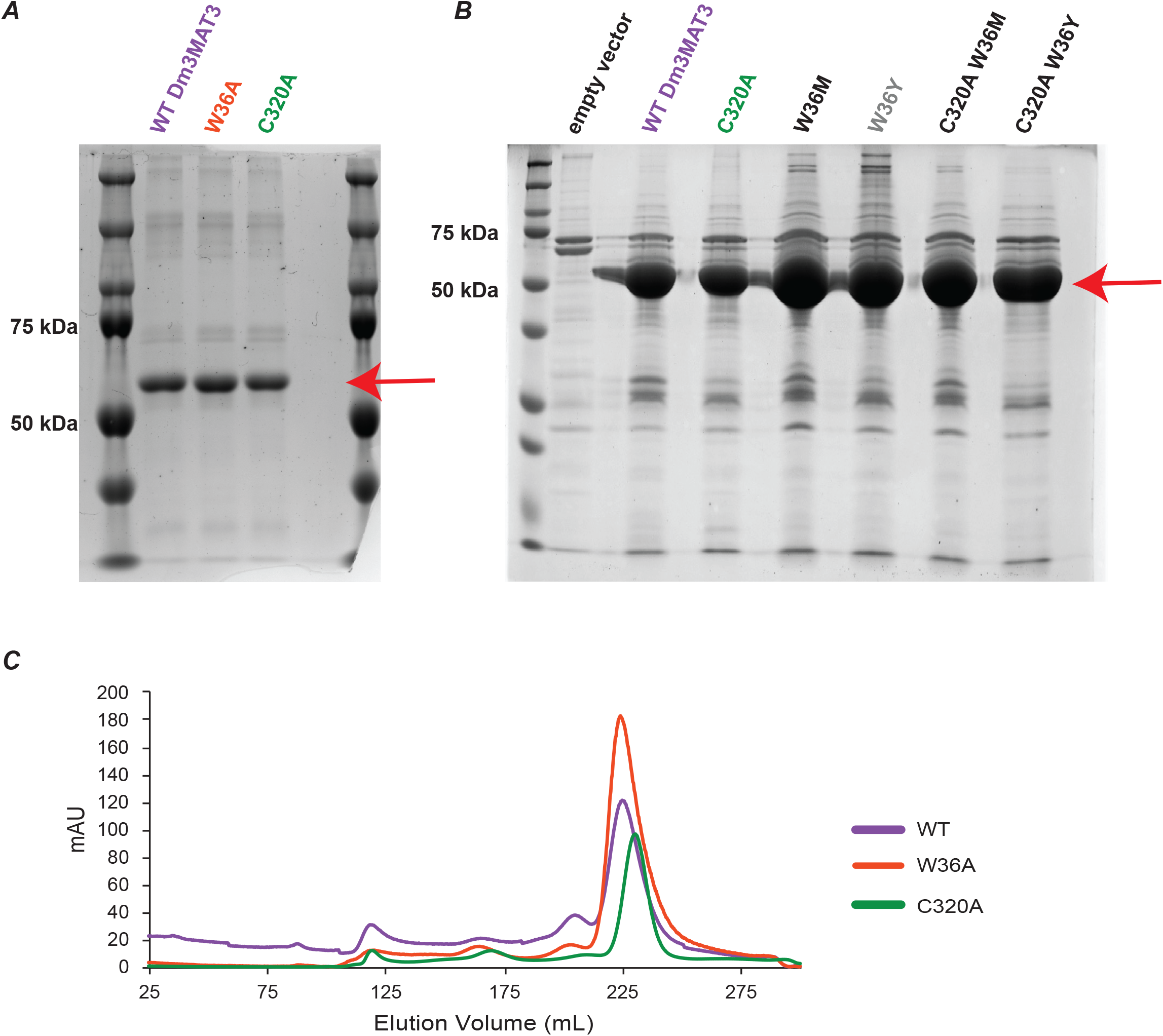
SDS-PAGE and Size Exclusion Chromatography of Dm3MAT3 variants. **(A)** Coomassie-stained SDS-PAGE of Dm3MAT3 variants purified to homogeneity using Ni-NTA-agarose. Similar migration pattern and purity for all variants was observed. **(B)** Coomassie-stained SDS-PAGE of further Dm3MAT3 variants purified to homogeneity using Ni-NTA-agarose. Similar migration pattern and purity for all variants was observed. **(C)** Size exclusion chromatography traces of the different Dm3MAT3 variants that have been used for kinetic measurements show that mutant proteins retain to similar elution volumes as wild-type, indicating proper size and shape of the protein (**Supp. Dataset 9**). Red arrow indicates expected size for Dm3MAT3 and its variants accoding to the protein ladder.

**Supplementary Figure 14.**
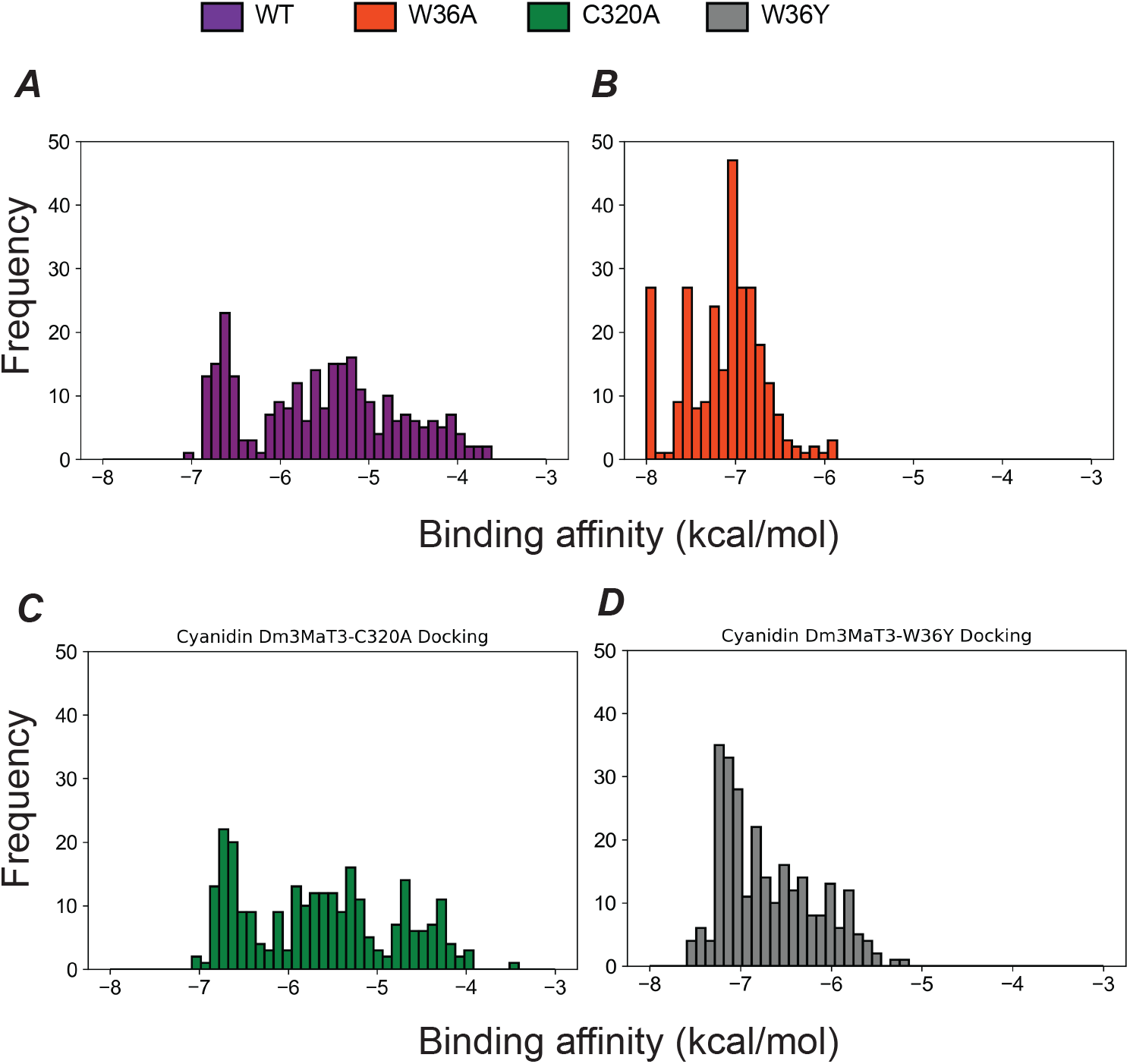
Substrate docking. Cyanidin 3-O-glucoside (C3G) docking into the Dm3MaT3 acceptor site for **(A)** WT, **(B)** W36A, **(C)** C320A, and **(D)** W36Y variants. WT is represented as purple, W36A is represented as orange-red, C320A is represented as green, and W36Y is represented as dark gray.

